# Identification of cell types in multiplexed *in situ* images by combining protein expression and spatial information using CELESTA reveals novel spatial biology

**DOI:** 10.1101/2022.02.02.478888

**Authors:** Weiruo Zhang, Irene Li, Nathan E. Reticker-Flynn, Zinaida Good, Serena Chang, Nikolay Samusik, Saumyaa Saumyaa, Yuanyuan Li, Xin Zhou, Rachel Liang, Christina S. Kong, Quynh-Thu Le, Andrew J. Gentles, John B. Sunwoo, Garry P. Nolan, Edgar G. Engleman, Sylvia K. Plevritis

**Affiliations:** Department of Biomedical Data Science, School of Medicine, Stanford University, CA, USA; Department of Radiology, School of Medicine, Stanford University, CA, USA; Cancer Biology Program, School of Medicine, Stanford University, CA, USA; Department of Pathology, School of Medicine, Stanford University, CA, USA; Stanford Cancer Institute, Stanford University, CA, USA; Division of Head and Neck Surgery, Department of Otolaryngology, School of Medicine, Stanford University, CA, USA; Department of Computer Science, Stanford University, CA, USA; Department of Radiation Oncology, School of Medicine, Stanford University, CA, USA; Department of Medicine, Stanford University, CA, USA

## Abstract

Advances in multiplexed *in situ* imaging are revealing important insights in spatial biology. However, cell type identification remains a major challenge in imaging analysis, with most existing methods involving substantial manual assessment and subjective decisions for thousands of cells. We propose a novel machine learning algorithm, CELESTA, which uses both cell’s protein expression and spatial information to identify cell type of individual cells. We demonstrate the performance of CELESTA on multiplexed immunofluorescence *in situ* images of colorectal cancer and head and neck cancer. Using the cell types identified by CELESTA, we identify tissue architecture associated with lymph node metastasis in HNSCC, which we validate in an independent cohort. By coupling our *in situ* spatial analysis with single-cell RNA-sequencing data on proximal sections of the same tissue specimens, we identify and validate cell-cell crosstalk associated with lymph node metastasis, demonstrating the power of spatial biology to reveal clinically-relevant cellular interactions.

## INTRODUCTION

Spatial biology is a new frontier that provides unprecedented characterization of tissue architectures through technological advances in multiplexed *in situ* imaging platforms ^1–6^. Using these platforms, specific tissue architectures have been associated with tissue development, disease progression and treatment response ^7–10^. However, such insights often rely on substantial data preprocessing that can introduce biases. In the case of *in situ* image, raw pixel data often need to be converted into images of cells through cell segmentation followed by identification of cell types for individual cells. Current cell type identification methods typically involves manual gating and/or clustering cells based on their protein expression profiles. Moreover, manual gating is subjective ^11,12^ and becomes unmanageable with high dimensional data. Clustering cells with similar protein expressions can be biased through the selection of number of clusters and it is often the case that individual clusters are a mixture of cell types, thereby comprising the single-cell resolution of the data ^13^. Even after clustering, decisions to assign cell types to the clusters are subjective, and clusters with cell type mixtures particularly demand much manual assessment. Given these limitations, downstream analysis of cellular spatial patterns can be highly compromised.

To address the limitations of common cell type identification methods, we reason that cells are organized in coherent spatial patterns and a cell’s spatial neighborhood is valuable information and could be considered together with cell’s protein expression profile to infer its cell type. We propose a novel machine learning cell type identification method, termed “CELESTA” (CELl typE identification with SpaTiAl information), that incorporates both cell’s protein expression profile and its spatial information, with minimal to no user-dependence, to produce relatively fast cell type assignments. CELESTA assigns cells to their most likely cell types through an optimization framework based on Markov Random Field modeling and relies on an input of prior information on cell type markers, thereby leveraging biological knowledge in a non-subjective manner.

We demonstrate the performance of CELESTA on imaging data generated using the CODEX (CO-Detection by indEXing) platform ^14^. CODEX is an immunofluorescence-based multiplexed *in situ* imaging technology that quantifies a cell’s spatial location and expressions of over fifty proteins, across tens of thousands of cells of a tissue slice ^14^. We applied CELESTA to a published CODEX dataset generated on colorectal cancer where cell type identification is based on both unsupervised and supervised clustering and manual assessment by pathologists ^6^, which we adopt as the gold standard for benchmarking CELESTA. CELESTA provides cell type assignments comparable to the gold standard. Moreover, CELESTA provides assignments to cells grouped in clusters which were left as unassessed through manual annotation because the clusters were regarded a mixture of different cell types. Since CELESTA does not rely on clustering, it was able to identify these unlabeled cell types.

To identify tissue architectures associated with lymph node metastasis, we applied CELESTA on our study cohort of head and neck squamous cell carcinoma (HNSCC) where we generated CODEX images from patients with (N+) and without (N0) lymph node metastasis. We identified specific cell types that were hypothetically colocalized only in N+ HNSCC patients and validated our findings on an external HNSCC cohort of tissue microarray (TMA). By coupling our spatial analysis with single-cell RNA-sequencing data on proximal sections of the same tissue specimens of the imaging data, we identified and validated cell-cell crosstalk associated with lymph node metastasis, demonstrating the power of spatial biology to reveal clinically-relevant cell-cell interactions.

## RESULTS

### Analysis Overview

Our analysis pipeline was designed to associate tissue architecture in a primary tumor with the nodal status (**Figure 1a**). Our study started with multiplexed *in situ* images on which we performed segmenting pixel-based data into cells, and identified cell types for each segmented cell using CELESTA. We then performed spatial analysis to characterize the tissue architecture to identify statistically-significant cell-cell colocalization patterns associated with nodal status and generate spatial biological hypothesis. By coupling this spatial biology with scRNA-seq analysis, we identified potential cell-cell crosstalk associated with lymph node metastasis.

**Figure 1.**
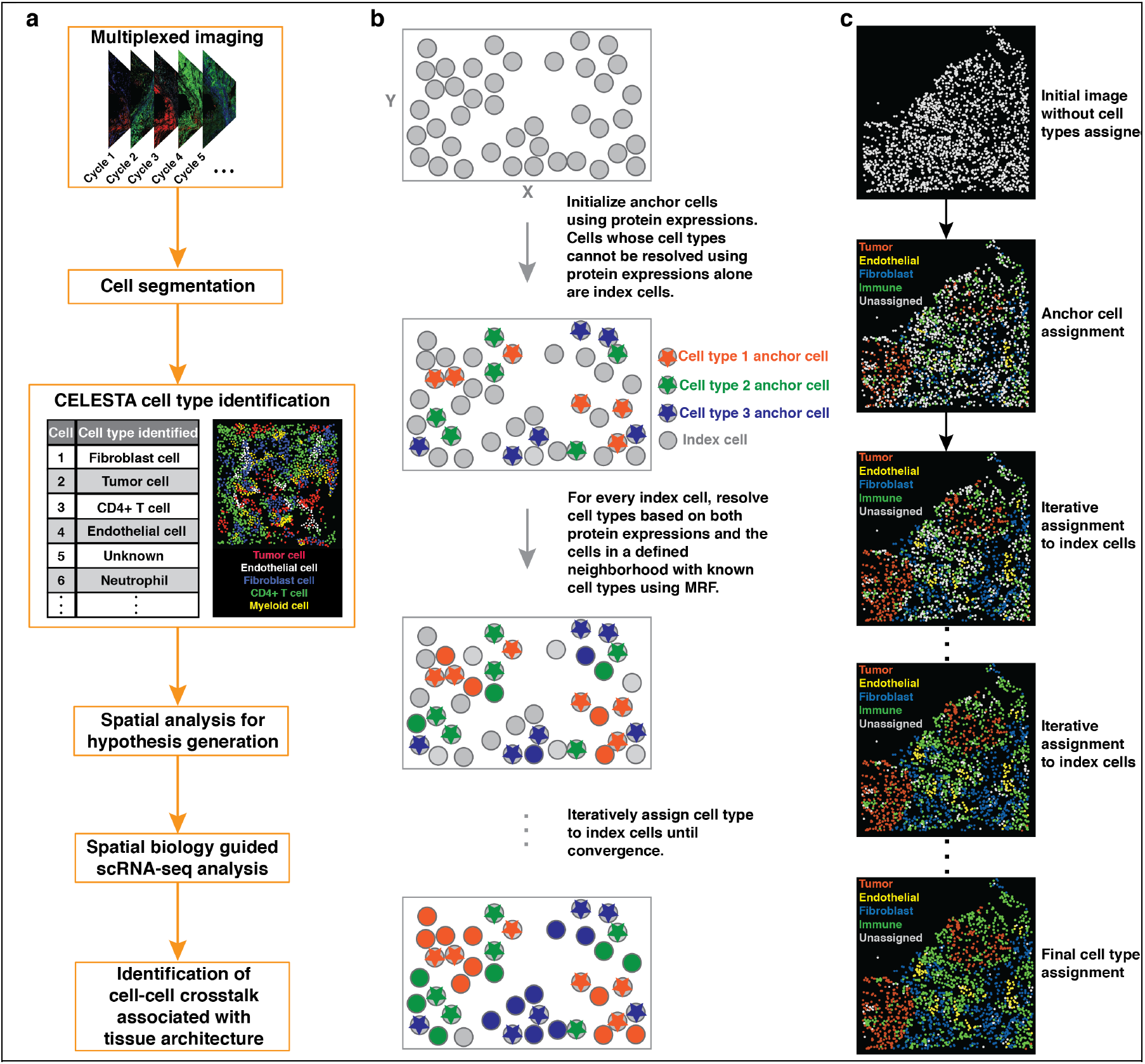
Study pipeline and overview of CELESTA. **(a)** A multiplexed *in situ* imaging data analysis pipeline used in the study. CELESTA identified cell types can be used to generate hypothesis of tissue architecture and guide scRNA-seq analysis to discover cell-cell crosstalk mediating tissue architecture. **(b)** Schematic illustration demonstrates CELESTA cell type assignment process in the tissue. MRF: Markov Random Field. **(c)** Illustration of CELESTA on a real image tile from a tissue sample.

### Overview of CELESTA

CELESTA is an unsupervised machine learning cell type identification method. CELESTA is currently optimized for *in situ* multiplexed immunofluorescence-based imaging but has broader applicability. Post cell segmentation, CELESTA first assigns cell types to cells whose protein expression profiles clearly match prior knowledge of cell-type-specific markers; those cells are defined as “anchor cells.” For the remaining cells, whose protein expression profile does not have clear pattern associated with a cell type, referred to hereon as “index cells”, CELESTA uses the neighboring cell-type information in addition to the cell’s protein expression profile to identify the cell type. Cell types for index cells are assigned through an iterative optimization framework. Illustration of CELESTA cell type assignment in the tissue is provided in **Figure 1b and 1c.** A flowchart illustrating CELESTA algorithm is shown in **Figure 2a**.

**Figure 2.**
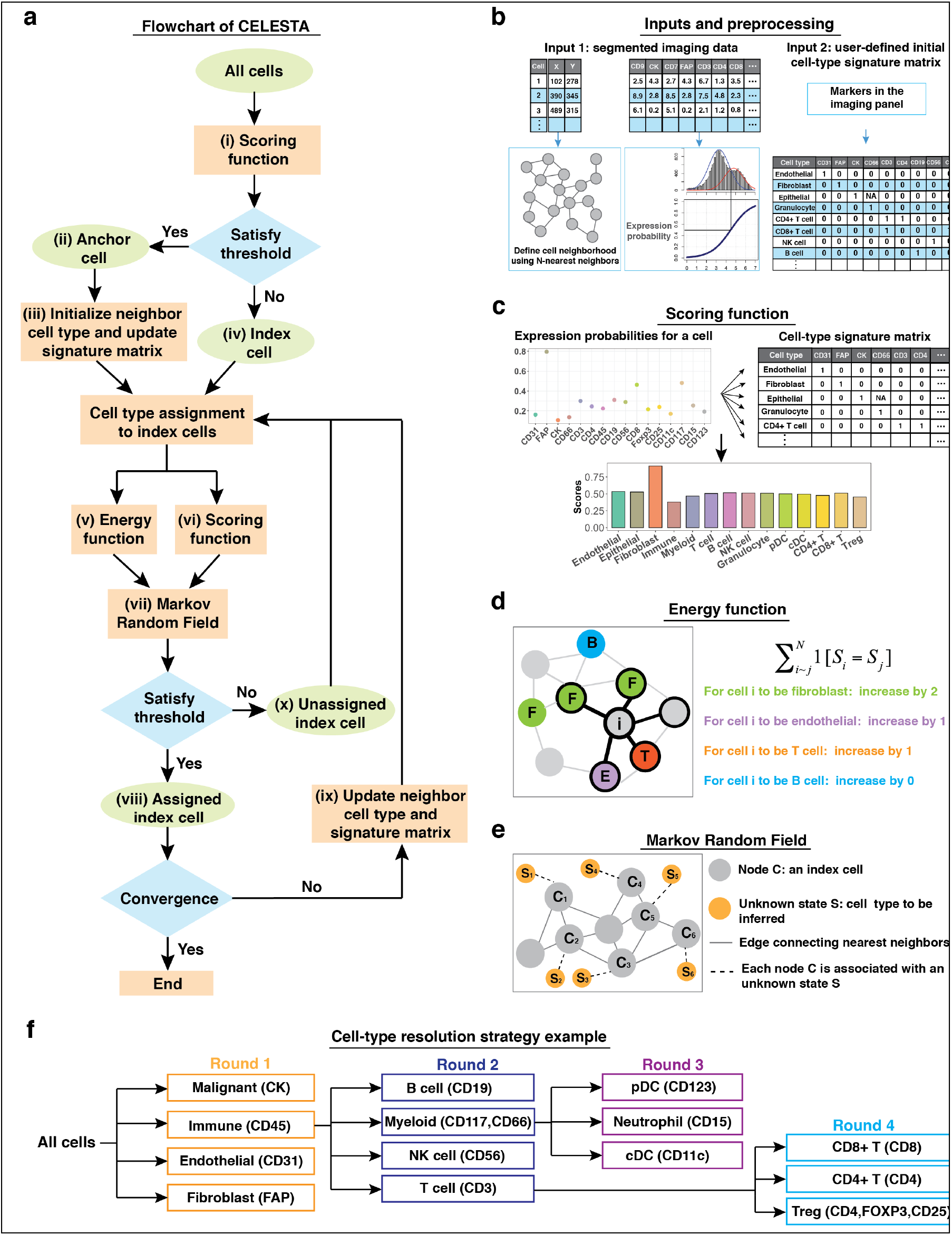
Illustration of key steps in CELESTA. **(a)** Flowchart of CELESTA algorithm. **(b)** CELESTA inputs and data processing. **(c)** An example illustrates CELESTA scoring unction. pDC: plasmacytoid dendritic cell. cDC: conventional dendritic cell. CK: cytokeratin. **(d)** An example illustrates neighborhood of an index cell *i*. The cell type information from spatial nearest neighboring cells of index cell *i* is added to the field by an energy function using Potts model indicator function. **(e)** We assume that each index cell *C* is associated with an unknown state *S* which is the cell type to be inferred. Cells are represented as nodes in an undirected graph with edges connecting the neighboring cells. We assume that the joint distribution of *S’s* satisfies discrete Markov Random Field. **(f)** An example demonstrating cell-type resolution strategy based on our HNSCC imaging panel.

CELESTA requires two inputs. The first input is an *in situ* image that has been segmented into individual cells where each cell is defined by its protein expression profile and spatial location, described in terms of X and Y coordinates (**Figure 2b**). In a preprocessing step, CELESTA defines the “cell neighborhood” using N-nearest neighbors based on cell’s spatial X and Y coordinates (**Figure 2b**). CELESTA then determines whether or not a protein marker is overexpressed in a given cell relative to other cells by fitting a two-mode Gaussian mixture model to the protein expressions across all the cells in a given sample ^15^; this step identifies the distributions for high vs. low expression levels. CELESTA then converts the protein expression into an “expression probability” based on a sigmoid function where the midpoint is the intersection of posterior probabilities based on the two-mode Gaussian distribution (**Figure 2b**). After preprocessing, each protein expression is scaled between 0 and 1 as a probability.

The second input to CELESTA is a cell-type signature matrix that defines each cell type in terms of its protein signature. This signature matrix relies on prior knowledge of proteins known to be expressed in specific cell types, given protein markers in the imaging panel (**Figure 2b**). For each cell type, the cell-type signature matrix indicates whether a protein is expressed in that cell type, initialized as 1 if expressed or 0 if not expressed, and updated with cells assigned. If the expression of a specific protein is irrelevant to identity the cell type, the protein is denoted as “NA” for that cell type. An example initial cell-type signature matrix on the public colorectal cancer dataset is shown in **Table S1**.

For cell type assignment, CELESTA first matches a cell’s expression probabilities to cell-type signatures to determine if it is sufficient to identify its cell-type (**Figure 2a(i)**). To make the assessment, CELESTA relies on a scoring function (**Figure 2a(i), 2c**), defined as 1 minus the mean squared errors between expression probabilities of a cell and the reference protein expression profile for each cell type in the initial cell-type signature matrix (**Figure 2c**). When a cell has one dominate cell type score, CELESTA assigns the corresponding cell type to that cell and defines the cell as an “anchor cell” (**Figure 2a(ii)**). CELESTA then updates the cells’ neighborhood cell types and cell-type signature matrix (**Figure 2a(iii)**) using the information from anchor cells.

For a cell whose cell type cannot be identified using protein expression alone (index cell, **Figure 2a(iv)**), CELESTA defines an energy function (**Figures 2a(v), 2d**) using the Potts model ^16^ to leverage cell type information on its spatially N-nearest neighboring cells in a probabilistic manner. The Potts model has been used for image segmentation where an indicator function is applied to increase the potential of the neighboring pixels belonging to the same object ^17–19^. The Potts model has also been demonstrated useful to analyze pathological images ^20^. CELESTA extends the concept of Potts model to spatial neighborhoods of cells, such that if there is one dominant cell type among the neighboring cells, that cell type contributes higher potential. Incorporating both energy function (**Figure 2a(v)**) and scoring function (**Figure 2a(vi)**), CELESTA represents each index cell as a node in an undirected graph with each edge connecting its spatially N-nearest neighbors. CELESTA associates each node with a hidden state, where the hidden state is the cell type to be inferred, and assumes that the joint distribution of the hidden states satisfy discrete Markov Random Field (**Figure 2a(vii), 2e**). To maximize the joint probability objective function, CELESTA employs an EM algorithm using mean field approximation ^21^. In each iteration, cell types with maximum clear-cut probabilities are assigned to the index cells (**Figure 2a(viii)**). The neighborhood cell types and signature matrix are also updated in each iteration (**Figure 2a(ix)**). If the protein expression profile together with the spatial information still does not produce clear-cut on probability for a cell type, CELESTA re-evaluates the cell on the next iteration with more neighboring cells assigned cell types (**Figure 2a(x)**). The process is repeated until remaining number of cells is smaller than a user defined convergence limit. The cells without cell type identities are assigned to an “unknown” category.

#### Incorporating cell lineage information into cell type identification

Because assigning cell types to tens of thousands cells is computationally expensive, CELESTA introduces a “cell-type resolution” strategy, whereby the cell type assignment is performed in multiple rounds where each round introduces increasing cell-type resolution based on known cell lineages **(Figure 2f)**. This approach not only reduces computational complexity but also improves robustness when cell types from different lineages have shared marker expression. More details on CELESTA are provided in Methods.

### Comparison of CELESTA with published annotations

We assessed the performance of CELESTA on a public domain CODEX dataset where we used the published cell types as “ground truth.” This dataset is a TMA composed 70 cores of human colorectal cancers. In the published study, the cell types were identified by applying both unsupervised ^22^ and supervised clustering together with pathologists’ manual assessment of marker expressions, cell morphology and staining colocalization ^6^. The CELESTA assignments, which were automated and computed on the order of minutes, were consistent with expert assessment. A comparison of CELESTA and published annotations on a representative TMA core is shown in **Figure 3a**. The number of cells identified per TMA core for each cell type were highly correlated between CELESTA and published annotations (**Figure 3b,c**). Using the published cell type assignments as the gold-standard, we showed that CELESTA achieved an average accuracy score (Rand Index) of around 0.9 across the cell types (**Figure 3d**); CELESTA has average precision scores ranged between 0.6 to 0.8 and F1 scores ranged between 0.6 to 0.7 across the major cell types (**Figure 3d**). For rare populations that are hard to identify, CELESTA achieved average precision and F1 scores ranged between 0.4 to 0.6. Noteworthy, there are two clusters in the published annotations that were assigned as cell type mixtures (**Figure 3e, f**); for cells in these two clusters, CELESTA assigned cell types matched with canonical marker expression patterns (**Figure 3e, f**), demonstrating CELESTA’s unique ability to provide cell type identification at single-cell resolution, not relying on clustering.

**Figure 3.**
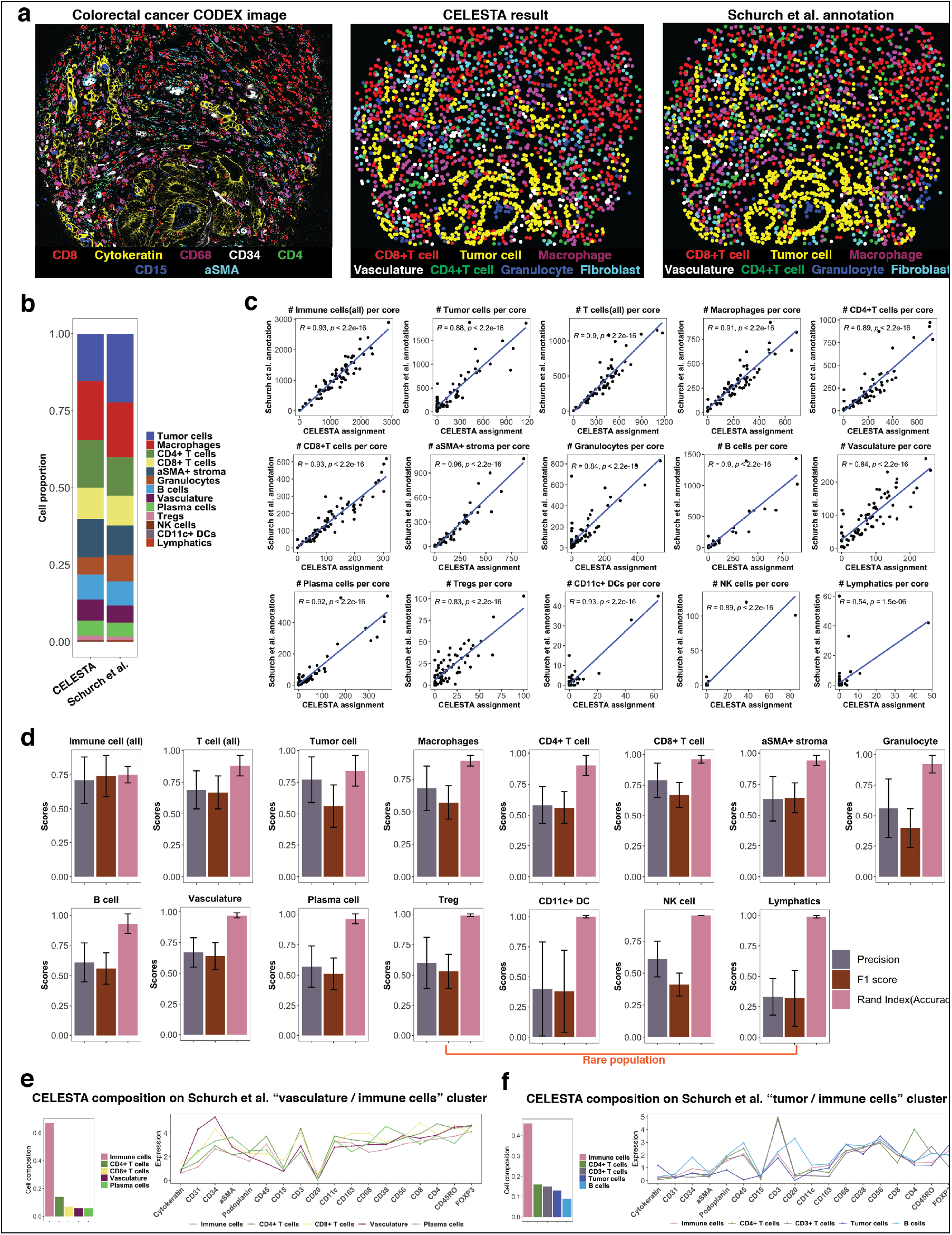
CELESTA applied to a public CODEX dataset of colorectal cancer tissue microarray (Schurch et al.). **(a)** An example tissue microarray (TMA) core region of CODEX image with seven channel overlay (Left), image derived from CELESTA assigned cell types (Middle) and image derived from Schurch et al. annotated cell types (Right). **(b)** Cell type compositions from CELESTA inferred cell types and Schurch et al. annotations. **(c)** Correlation diagrams show number of cells identified per TMA core across 70 core regions between CELESTA and Schurch et al. annotations for each cell type. Numbers of cells identified per TMA core region for each cell type are correlated between CELESTA identified cell types and Schurch et al. annotations. **(d)** Precision scores, F1 scores, and accuracy (Rand Index) scores for major cell types identified by CELESTA using Schurch et al. annotations as “ground truth”. Error bars are calculated using standard deviations across 70 TMA core regions in the data. Rare population is defined as average of fewer than 20 cells per core region in the annotations. **(e)** CELESTA cell type assignments on a cluster of cells which Schurch et al. annotated as a mixture of vasculature or immune cells. CELESTA cell type compositions are shown on the left panel and average canonical marker expressions for each cell type in the cluster are shown on the right panel. **(f)** CELESTA cell type assignments on a cluster of cells which Schurch et al. annotated as a mixture of tumor or immune cells. CELESTA cell type compositions are shown on the left panel and average canonical marker expressions for each cell type in the cluster are shown on the right panel.

To evaluate the mismatched cell type assignments between CELESTA and published assignments, we evaluated the protein expression profiles. For example, among the tumor cells that we mismatched, we analyzed the protein expressions of the tumor cells identified by: (i) both methods, (ii) only in the published annotations and (iii) only by CELESTA (**Figure S1**). Tumor cells that were identified the published annotations but not by CELESTA express very low to none cytokeratin, which is a common tumor cell marker. Tumor cells identified by CELESTA express cytokeratin as defined in the input of cell-type signature matrix.

### CELESTA applied to primary HNSCC tumors imaged by CODEX

We generated a study cohort of eight primary HNSCC tumors, comprised of four node-positive (N+) and four node-negative (N0) samples (**Table S2**). We performed CODEX imaging using the protein marker panel in **Table S3**. The input cell-type signature matrix based on the imaging panel is shown in **Table S4**. We manually assessed CELESTA’s performance by mapping CELESTA assigned cell types onto the CODEX images using the X and Y coordinates against the canonical marker staining (Methods). We showed qualitatively that CELESTA assigned cell types matched well with canonical marker staining (**Figure 4a, 4b, S2**). Next, we evaluated cell type composition from CELESTA with paired scRNA-seq data on proximal tissue sections from four samples in our HNSCC cohort (two N+ and two N0 samples). CELESTA identified cell type compositions were correlated with scRNA-seq cell type compositions (R=0.7, P<0.0001, **Figure 4c**). While the cell type compositions are concordance, the differences may arise because tissue dissociation methods used for generating scRNA-seq data often over-enriched for immune cells.

**Figure 4.**
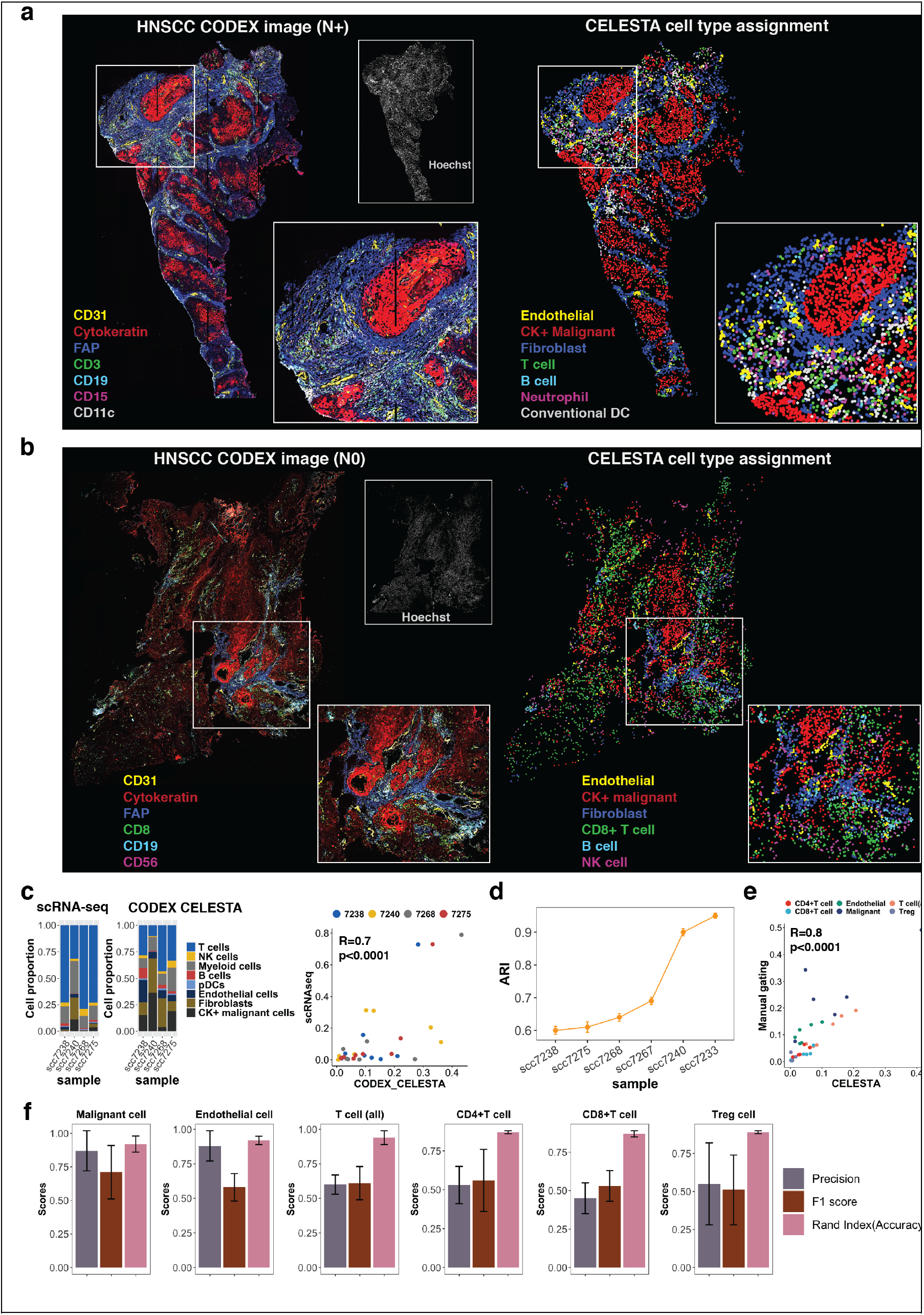
CELESTA applied to CODEX data of fresh frozen HNSCC primary tumor samples. **(a)** CODEX image overlay and CELESTA results on major cell types in a primary tumor tissue sample of head and neck squamous cell carcinoma (HNSCC) with lymph node metastasis (N+). CK: Cytokeratin. **(b)** CODEX image overlay and CELESTA results on major cell types in a primary tumor tissue sample of HNSCC without lymph node metastasis (N0). **(c)** Cell type compositions from scRNA-seq data (Left) and CELESTA inferred cell types on CODEX data (Middle). scRNA-seq data were obtained from proximal tissue slices with the CODEX data tissue on the same four patient samples. Correlation between CELESTA inferred cell compositions and scRNA-seq cell compositions on the same four samples (Right). **(d)** Adjusted rand index (ARI) to assess CELESTA’s performance against manual gating for each tissue sample. Error bars were calculated based on 50 runs of random sampling. **(e)** Correlation between CELESTA inferred cell compositions and manual gating compositions. **(f)** Cell type precision scores, F1 scores and accuracy scores (Rand index) on cell types relevant to downstream analysis (malignant cells, endothelial cells and T cell subtypes).

To further quantitatively evaluate the performance of CELESTA, we applied manual gating as benchmark. Due to the limitation of gating on high dimensionality data, we designed gating strategies focusing on cell types relevant with downstream analysis (Methods and **Figure S3**). Compared with manual gating, CELESTA achieved sample adjusted rand index (ARI) ranged from 0.6 to 0.9 across the samples (**Figure 4d**). Due to imaging artifacts, cell type identification was harder in some tissue samples. In terms of cell type compositions, CELESTA and manual gating were highly correlated (R=0.8, p<0.0001, **Figure 4e**). For cell-type specific assessment, CELESTA achieved average F1 scores of around 0.7 and average accuracy scores of around 0.9 for malignant cells, endothelial cells and T cells (**Figure 4f**). For T cell subtypes, CELESTA achieved average F1 scores around 0.55 and average accuracy scores around 0.9 compared with manual gating (**Figure 4f**). Because T cell subtypes were identified in later rounds of manual gating sequence, potential accrual manual errors could cause reduced accuracy measures.

### CELESTA-enabled spatial analysis reveals differential cell-cell co-localization patterns in primary HNSCC dependent on nodal status

We performed spatial analysis on our HSNCC study cohort using the cell types identified by CELESTA. We adapted the co-location quotient ^23^ (CLQ), a measurement used in geospatial statistics, to quantify spatial co-localization between pairs of cell types and test the hypothesis that there are differential cell-type-specific co-localization patterns in N+ vs. N0 HNSCC (**Figure 5a**). Specifically, by denoting cell-type *a* as target cells and cell-type *b* as neighboring cells, we used CLQ to measure the degree to which cell-type *b* is spatially co-localized with cell-type *a* and calculated the CLQs for each pair-wise cell types for each sample and compared the CLQs between N+ and N0 samples. Four pairs of cell types that were significantly more co-localized in N+ vs. N0 samples (**Figure 5b**) (p-values < 0.05), namely: (i) Malignant cells and Treg cells, (ii) CD4+ T cells and endothelial cells, (iii) CD8+ T cells and CD4+ T cells, and (iv) CD4+ T cells with themselves. The hypothetical spatial co-localization pattern differences between N0 and N+ samples are illustrated in **Figure 5c**. Representative pairs of CODEX images show FOXP3 (a canonical Treg marker) expression is more co-localized with cytokeratin (tumor marker) expression in N+ vs. N0 HNSCC (**Figure 5d**), and CD4 and CD8 (canonical T cell markers) expression are more co-localized with CD31 (a canonical endothelial marker) expression in N+ vs. N0 HNSCC (**Figure 5e**). To validate the hypothesis of co-localization of Tregs and malignant cells in N+ HNSCC, we stained FOXP3 and cytokeratin on a TMA from an independent cohort of 76 HNSCC patients. Representative TMA images illustrating more co-localization of cytokeratin-positive malignant cells and Tregs in N+ vs. N0 samples are provided in **Figure 5f.** We showed that quantitatively primary N+ HNSCC have significantly higher density correlations between cytokeratin and FOXP3 (p < 0.05, **Figure 5g**) using a TMA-image processing pipeline (Methods).

**Figure 5.**
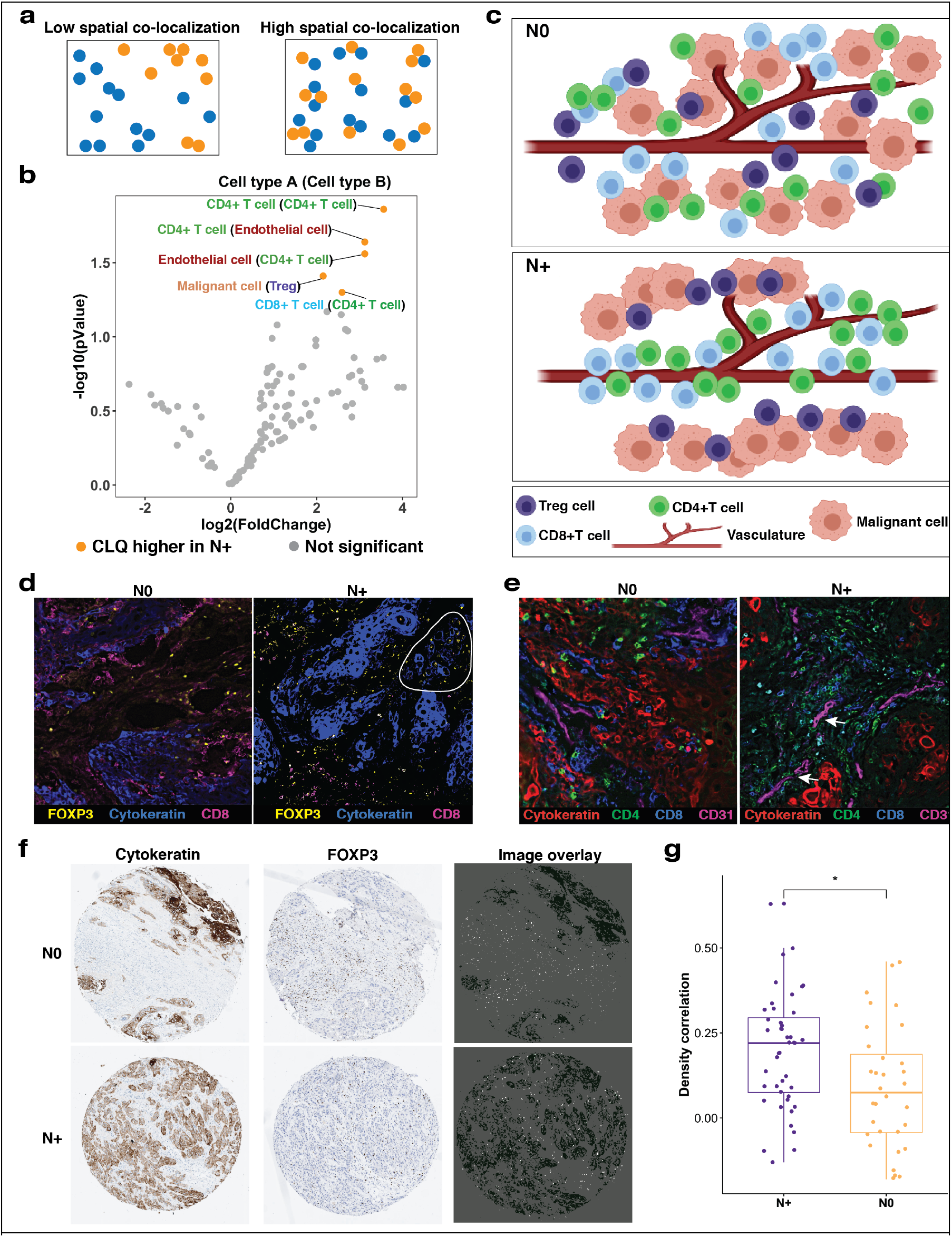
Spatial cell-cell co-localization analysis based on CELESTA identified cell types in the HNSCC study cohort. **(a)** Schematic representation of two different cell spatial patterns respectively for low cell co-localization (Left) and high cell co-localization (Right). **(b)** Comparing co-localization quotients (CLQs) for pair-wise cell types between N+ and N0 HNSCC samples. **(c)** Schematic illustrations of hypothetical tissue spatial architectural differences of cell-cell co-localizations in N0 samples (Top) and N+ samples (Bottom). **(d)** Representative regions for N0 sample (Left) and N+ sample (Right) depicted as three-color overlay images with FOXP3 (yellow), Cytokeratin (blue) and CD8 (magenta). **(e)** Representative regions for N0 sample (Left) and N+ sample (Right) depicted as four-color overlay images with Cytokeratin (red), CD4 (green), CD8 (blue) and CD31 (magenta). **(f)** Representative HNSCC TMA cores for N0 and N+ patients depicted as overlay images with Cytokeratin and FOXP3 staining. **(g)** Density correlation analysis shows that Cytokeratin and Foxp3 have higher density correlations in N+ patients (n=44) than N0 patients (n=32) in an independent TMA cohort of HNSCC samples. *p-value < 0.05

### Spatially-guided scRNA-seq analysis implicates cell-cell crosstalk mediating tissue architecture associated with lymph node metastasis in HNSCC

By reasoning that crosstalk between distinct cell types may be associated with physical proximity ^24,25^, we sought spatial co-localization patterns as a guide to discover cell-cell crosstalk mediators associated with nodal status. We leveraged our HNSCC atlas, derived from scRNA-seq data that was generated on tissue specimens proximal to the imaged specimens (**Figure 6a, S5a**) using Seurat ^26,27^. In this atlas, we identified a malignant cell cluster (Cluster 11) in which CXCL10, an interferon-inducible chemokine ligand, was more expressed on N+ vs. N0 HNSCC (**Figure 6b, 6c**). We also identified a T cell cluster (Cluster 2) as Treg-enriched, based on FOXP3 and IL2RA expression (**Figure S5b**), and found that CXCR3, a receptor of CXCL10, was more expressed on N+ vs. N0 HNSCC in the Treg cluster (**Figure 6c**). This finding led us to hypothesize the CXCL10-CXCR3 interaction as an axis for crosstalk between malignant cells and Tregs in N+ HNSCC. Evidence for this CXCL10-CXCR3 interaction was found in an independent public scRNA-seq HNSCC dataset ^28^ (**Figure S6**). In a similar manner, we found that the ligand-receptor pair CCL20-CCR6 is higher in N+ vs. N0, and may mediate crosstalk between endothelial and CD4+ T cells in N+ HNSCC (**Figure 6d**). In summary, we demonstrate that CELESTA enabled cell-cell co-localization patterns that guided scRNA-seq analysis to identify cell-cell crosstalk mediators related to lymph node metastasis (**Figure 6e**).

**Figure 6.**
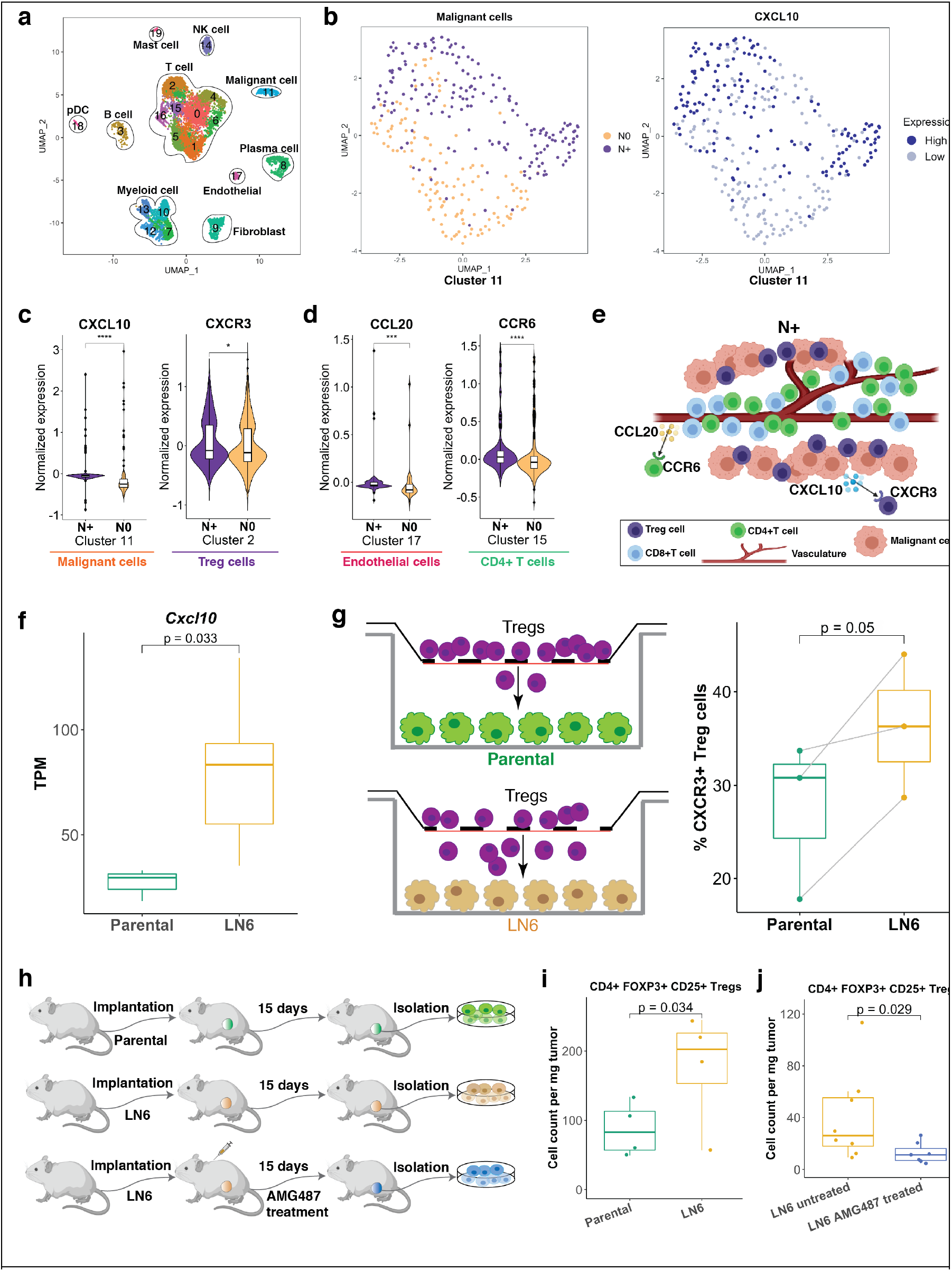
Spatial biology guided scRNA-seq analysis. **(a)** UMAP plot of identified cell type clusters using paired HNSCC scRNA-seq data. **(b)** UMAP plots of malignant cells (Cluster 11) with node status (Left) and CXCL10 expression (Right). **(c)** Violin plots show CXCL10 and CXCR3 expressions are differentially expressed in malignant cell and Treg cell clusters respectively comparing N+ vs. N0 samples. **(d)** Violin plots show CCL20 expressions in endothelial cell cluster and CCR6 expressions in CD4+T cell cluster between N+ and N0 samples. **(e)** Schematic illustration shows cell-cell crosstalk with identified chemokine ligand-receptor pairs mediating cellular spatial co-localization in N+ samples. **(f)** Cxcl10 expression is significantly higher in mouse model LN6 generation cell lines than parental cell lines. **(g)** Transwell experiment shows that LN6 malignant cells attract more CXCR3+ Treg cells through the membrane than parental cell lines. **(h)** Schematic workflow of *in vivo* experiments. **(i)** LN6 line tumors recruit more Treg cells into the tumors than parental tumors. **(j)** AMG487 treatment significantly reduces Tregs recruited into the LN6 tumors. *: adjusted p-value < 0.05, **: adjusted p-value < 0.01, ***: adjusted p-value < 0.005, ****: adjusted p-value < 0.001.

### Functional validation on CXCL10-CXCR3 crosstalk between malignant cells and Tregs

To validate the association of CXCL10-CXCR3 between malignant cells and Tregs with nodal status, we leveraged an *in vivo* model of lymph node (LN) metastasis ^29^ that we developed in melanoma. In this model, we created multiple generations of lymph node metastatic cell lines (LN1-LN6), with each generation exhibiting an increased frequency of lymph node metastases. RNA sequencing of different LN lines revealed that later generations (LN6) expressed significantly higher levels of CXCL10 compared with the parental cell line (**Figure 6f**), indicating metastatic cells to the lymph node upregulate CXCL10.

We tested the hypothesis that CXCR3+ Tregs are more attracted to CXCL10+ malignant cells in a transwell experiment. Tregs harvested from naive FOXP3-GFP mice were plated in the upper chambers; the bottom chambers were plated with either the parental melanoma cells (control group) or LN6 melanoma cells (study group). We found that LN6 cells induced more migration of CXCR3+ Treg cells through the membrane compared to parental cells (**Figure 6g**). With higher expression of CXCL10 and higher frequency of lymph node metastasis in the LN6 line compared with parental line, this finding is in accordance with our hypothesis that CXCR3-CXCL10 promotes Treg migration toward malignant cells that colonize lymph nodes.

Given the existence of a small molecular weight antagonist, AMG487, that has shown to block CXCR3 and reduce metastasis ^30–32^, we hypothesized that Treg tumor infiltration is
reduced following AMG487 treatment. We created an *in vivo* experiment to compare the migration of Treg cells into parental tumor versus LN6 tumor with and without AMG487 treatment (**Figure 6h**). We found that LN6 tumors recruited more Tregs than the parental line tumors *in vivo* (**Figure 6i**). Following AMG487 treatment on LN6 tumors, the number of Tregs recruited into the tumor was reduced (**Figure 6j**), validating our hypothesis.

## DISCUSSION

Spatial biology is a new frontier that has become accessible through advances in multiplexed *in situ* imaging. Exploring this frontier involves converting raw pixel-based images into an interpretable cell-based format. This poses numerous technical challenges, among them is the identification of cell types of individual cells. Common strategies for cell type identification rely on manual gating or clustering algorithms. Gating and clustering require substantial manual assessment, making them largely subjective, time consuming and compromise single-cell resolution. We propose CELESTA, a novel machine learning method tailored for identifying cell types on highly multiplexed immunofluorescent *in situ* images by utilizing both the cell’s protein expression profile and spatial information. CELESTA is automated and fast (~10min for ~100k cells on a typical Macbook).

CELESTA has several important features. CELESTA’s input includes a cell-type signature matrix of user-defined prior knowledge of the cell types, and a scoring function that transfers this prior biological knowledge into cell type assignment in a non-subjective manner. CELESTA applies Gaussian mixture modeling to quantify whether a specific marker is expressed or not in a cell, relative to the expression in all other cells; this step relies on manual and subjective assessment in most existing cell type identification methods. In the situation when the cell’s protein expression profile is insufficient to determine the cell type, CELESTA incorporates cell’s spatial information to identify the cell type, which has not been considered by existing methods. CELESTA assigns the cell type to individual cells based on probabilities, and thus preserves the single-cell resolution of the data. Embedded in CELESTA a “cell-type resolution” strategy identifies cell types in multiple rounds based on cell lineage; this strategy not only improves computational speed but also improves robustness when cell types from different lineages have shared marker expression. CELESTA is able to handle large dataset with fast assessment of the image. We have shown that CELESTA has comparable performance against pathologist’s manual assessment. For rare cell types that are hard to identify, CELESTA’s probabilities provide extra information for further manual assessment. Lastly, we applied CELESTA to *in situ* images generated on a multiplexed immunofluorescence-based platform, but CELESTA could also be extended to data generated on other *in situ* imaging platforms.

Like other cell type identification methods, CELESTA requires segmenting cells in the images as input and thus it relies on the performance of segmentation algorithm which can be challenging with multiplexed imaging data. Particularly in the case of over-segmentation, CELESTA may over-weight the spatial neighborhood information. Tissue sample compositions and technical artifacts introduced by the imaging platform could add noise to the protein expression on some samples ^33,34^. In such cases, some manual intervention is still needed to obtain more accurate cell type identification after CELESTA’s fast assessment. CELESTA also relies on the cell-type marker information in the user-defined input cell-type signature matrix. A poorly informed cell-type signature matrix will effect the results. CELESTA, in its current version, does not account for cell size or cell morphology, which are also important information for cell type identification.

We showed that CELESTA-informed geospatial tissue analysis revealed novel spatial biology of HNSCC associated with lymph node metastasis, a finding which was validated on an external cohort. We also reasoned that crosstalk between distinct cell types may be associated with physical proximity ^24,25^ and thereby identified spatial co-localization patterns can provide guidance to discover potential cell-cell crosstalk mediators associated with nodal status. Using spatially-guided scRNA-seq analysis, we identified that the chemokine ligand-receptor pair CXCL10-CXCR3 have higher expressions on malignant cells and Tregs respectively in N+ vs. N0 HNSCC. While CXCL10-CXCR3 crosstalk has been suggested to play a critical role in T cell trafficking and cancer metastasis ^35–37^, our work provides a potential role of CXCL10-CXCR3 axis in mediating tissue architecture related to HNSCC lymph node metastasis. Using a small molecule weight antagonist of CXCR3 that reduced Treg tumor infiltration, we demonstrate the potential clinical importance of the CXCL10-CXCR3 axis as a therapeutic target for metastatic HNSCC. In a similar manner we also identified the CCR6-CCL20 axis, which was suggested to be related to cancer progression ^38–40^, mediating immune-endothelial crosstalk in node-positive disease.

In summary, we propose CELESTA as an automated and fast cell type identification method to facilitate multiplexed *in situ* image analysis, using both cells’ protein expression and spatial information. CELESTA enabled us to perform geospatial analysis and identify important spatial biology associated with nodal status in HNSCC. Our work also demonstrates the power of spatial biology to guide the discovery of cell-cell interactions associated with disease progression and thereby provide new therapeutic avenues.

## METHODS

### CELESTA

#### Scoring function

The first component in the algorithm is the scoring function to assess how well a cell’s marker expression profile match with our prior knowledge on the cell-type markers. To apply the scoring function, we first need to quantify whether a marker is expressed or not in a cell, which is a step often based on subjective assessment in most existing cell type identification methods. We apply a two-mode Gaussian mixture model to fit each marker’s expressions across all the cells ^15^ in a sample:

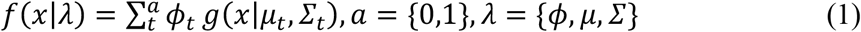

Where *ϕ* is the mixing probabilities that sum up to one, *μ* is the mean and *Σ* is the covariance. If we assume that an expressed marker is in state *a* = 1 and an unexpressed marker is in state *a* = 0, the posterior distributions for a marker to be expressed is *p*(*a* = 1|*x*) and unexpressed is *p*(*a* = 0|*x*). At the decision boundary, we have:

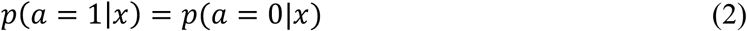

By using Bayes’ theorem, we obtain the following:

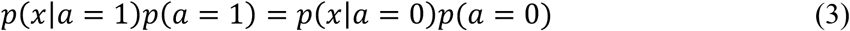

where *p*(*x*|*a* = 1) = *g*(*x*|*μ*_1_, *Σ*_1_) and *p*(*x*|*a* = 0) = *g*(*x*|*μ*_0_, *Σ*_0_). *p*(*a* = 1) and *p*(*a* = 0) are the mixing probability *ϕ*_1_ and *ϕ*_0_. All could be obtained from the Gaussian mixture model. By solving equation (3), we identify the decision critical point *x_c_* at which a marker has equal probabilities for expressing versus not expressing. We thus could use a logistic function to quantify a marker expression probability (*EP*) for each marker in each cell as:

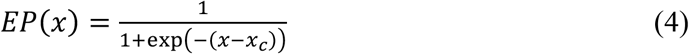

We repeat the process for every marker, and thus for each cell, its protein marker expressions are converted into expression probabilities. With the reference expression probabilities for markers expressed or not expressed for a cell type defined in the cell-type signature matrix for respective cell types, we define our scoring function for a cell type as one minus the mean squared error between each cell’s marker expression probabilities and the reference expressions in the cell-type signature matrix for that cell type. When a cell has one dominate cell type score, CELESTA assigns the corresponding cell type to that cell and defines the cell as an “anchor cell”. For a cell whose cell type cannot be identified using protein expression alone, it is defined as an index cell. In addition, a threshold for expressing and a threshold for not expressing based on the expression probability are also required. For example, we can set the high threshold to be 0.7 and low threshold to be 0.3 for tumor cells. This means that for a cell to be tumor cell, it needs to have cytokeratin (tumor marker) expression probability higher than 0.7 and all other non-expressing protein marker expression probability lower than 0.3, in addition to having a scoring function producing high score on tumor cell compare to other cell types. The thresholds provide user the flexibility to define artifacts like doublets or staining background noise. The cell-type signature matrix is updated in each iteration to incorporate the averaging expression probabilities from cells that are assigned cell types into the reference expressions.

CELESTA employs an optimization framework to assign the most likely cell type to each index cell. CELESTA is designed to maximize the joint probability distribution using hidden Markov Random Field (MRF) ^41^ that includes an energy function component that accounts for cell spatial information and a scoring function component that accounts for protein marker expressions. We assume each cell *C* whose cell type needs to be inferred (index cell) is a node in an undirected graph and each cell has connected neighboring cells that are stochastically dependent. We model the stochastic spatial dependency using neighboring system defined on the undirected graph *G* with the edges connecting the cell to its N-nearest neighboring cells. Depending on the cell densities in the tissue, we recommend N=5~10. We associate each node (an index cell) with an unknown state *S* which is the cell type to be inferred. Thus, the spatial dependency on the undirected graph is modeled by hidden Markov Random Field (MRF) with joint probability distribution of Gibbs distribution as:

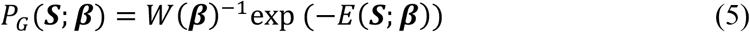

Where ***β*** is a set of model parameters to be estimated, *W*(***β***) is a normalization constant, and *E* is the energy function.

#### Energy function

For our energy function *E*, we use the Potts model ^17,18^, defined as:

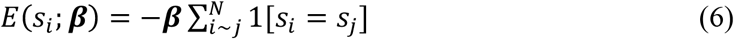

Where *N* is the number of nearest spatial neighboring cells of the index cell *i* based on cells’ X and Y coordinates obtained from the image. It is an indicator function. If an index cell has the same cell type as a neighbor cell *j*, one is added to the summation and thus increase the probability for that cell type. ***β*** is a set of model parameters depending on distances between cells. ***β*** is used to decide how much information to include from the neighboring cells, defined by triangular kernel times a user defined scale factor *γ* as:

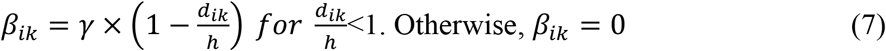

where *d_ik_* is the distance between index cell *i* and its nearest cell that has cell type *k* assigned, and *h* is the bandwidth. Therefore, if there are no cells of cell type *k* assigned within distance *h* to the index cell, no neighborhood information from cell type *k* is used. If an index cell is too far away from any cells with cell types identified, no spatial neighborhood information is used for that cell, which could help accounting for isolated cells. CELESTA uses the spatial dependency term to account for the situation where an index cell’s protein expression profile has similar scores matched to multiple cell types, reasoning that cells with the same cell type are enriched within each other’s spatial neighborhoods.

#### Optimization of objective function for cell type identification

With both the energy function and scoring function defined, our objective function is expressed as the overall joint probability distribution:

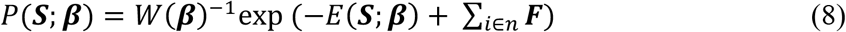

where *n* is the total number of index cells. *E* is the energy function component and *F* is the scoring function component. We intend to assign each individual cell with the cell types that maximize the joint probability distribution. Because our objective function is non-convex, we use a pseudo-EM algorithm to iteratively solve the problem. In the initialization step, we only use the scoring function to identify cells whose protein marker expressions could be well matched to only one cell type based on prior knowledge in cell-type signature matrix, and define those cells as anchor cells and assign the matched cell types to the anchor cells. Each iteration, we approximate the probability of cell type *k* for an index cell *i* using mean field approximation ^19^ by:

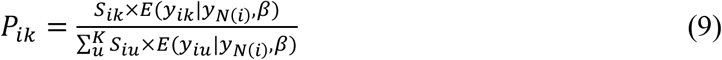

where *K* is total number of cell types in the cell-type signature matrix. If there are multiple rounds defined in the prior knowledge using “cell-type resolution” strategy, *K* is the total number of cell types in a round. *N* is the total number of neighbors of index cell *i*. Essentially, we approximate probabilities of each cell type for the index cell and assign the cell type with dominating probability to the index cell. A threshold of the probability is required. If the cell type probabilities do not pass the threshold, then no cell type is assigned for that cell in this current iteration and the cell is carried over to next iteration with more cell types assigned and increased neighborhood information. Each iteration, we update the parameters of *β*, the cell-type signature matrix, the scoring function and the energy function based on the new neighborhood information. The convergence condition is defined as the percentage of cells that do not have cell type assignment is smaller than a user-defined threshold. After the convergence condition is met, if a cell is still not assigned to a cell type, we assign the cell with an “Unknown” cell type.

### Human tumor specimens

All patients at the Stanford Hospital included were consented to take part in the study following Institutional Review Board (IRB) approval (IRB protocol #: 11402). The staging information of the patient samples are summarized in **Table S2**. Fresh tissue of head and neck squamous cell carcinoma (HNSCC) was collected within six hours of surgical resection. A two-to-three millimeter tissue piece was cut from the sample. Patient 7153, and 7155 OCT samples were immediately frozen in OCT freezing media, while patient 7233, 7238, 7240, 7267, 7268 and 7275 OCT samples were placed in 30% sucrose for 1 hour at 4° Celsius and frozen in OCT freezing media (Fisher Healthcare, Houston, TX) on a metal block chilled in liquid nitrogen. OCT samples were stored in - 80°C for CODEX processing and sequencing. The remaining tissue was placed on ice and in 50μl tissue digestion media, DMEM-F12+ with magnesium and calcium (Corning Cellgro, Manassas, VA), 1%FBS (heat inactivated), 10units/ml Penicillin-10ug/ml Streptomycin (Gibco, Grand Island, NY), 25mM hepes (Gibco, Grand Island, NY).

### CODEX image acquisition and segmentation

Multiplexed CODEX analysis of HNSCC tissues was performed using a panel of antibodies (**Table S3**) conjugated to custom DNA barcodes and detector oligos as well as common buffers, robotic imaging setup and instructions for CODEX staining of frozen specimen from Akoya Biosciences (https://www.akoyabio.com/). 7um sections were cut with a cryostat after OCT blocks were equilibrated to the cryostat temperature for at least 30-40 minutes. Tissue sections were placed on the surface of cold poly-L-lysine coated coverslips and adhered by touching a finger to the bottom surface to transiently warm up the coverslip. Frozen sections on coverslips can be stored at −70 C for 1-2 months. Section pre-processing was done as previously described ^14^. Prior to staining the sections, frozen sections removed from the freezer were dried for 5 min on the surface of Drierite. Dried coverslips with sections on them were dipped for 10 min into room temperature acetone, then fully dried for 10 min at room temperature. Sections were then rehydrated for 5 min in S1 [5 mM EDTA (Sigma), 0.5% w/v bovine serum albumin (BSA, Sigma)] and 0.02% w/v NaN3 (Sigma) in PBS (Thermo Fisher Scientific) then fixed for 20 min at room temperature in S1 with 1.6% formaldehyde. Formaldehyde was rinsed off twice with S1. Sections were equilibrated in S2 [61 mM NaH2PO4 · 7 H2O (Sigma)], 39 mM NaH2PO4 (Sigma) and 250 mM NaCl (Sigma) in a 1:0.7 v/v solution of S1 and doubly-distilled H2O (ddH2O); with final pH of 6.8-7.0 for 10 minutes, and placed in blocking buffer for 30 minutes. All steps to follow were exactly as previously described ^14^ or per the Akoya CODEX instructions.

Automated image acquisition and fluidics exchange were performed using an Akoya CODEX instrument driven by CODEX driver software (Akoya Biosciences) and Keyence BZ-X710 fluorescent microscope configured with 4 fluorescent channels (DAPI, FITC, Cy3, Cy5) and equipped with CFI Plan Apo λ 20x/0.75 objective (Nikon). Hoechst nuclear stain (1:3000 final concentration) was imaged in each cycle at an exposure time of 1/175s. Biotinylated CD39 - detection reagent was used at a dilution of 1:500, and visualized in the last imaging cycle using DNA streptavidin-PE (1:2500 final concentration). DRAQ5 nuclear stain (1:500 final concentration) was added and visualized in the last imaging cycle. Each tissue was imaged with a 20x objective in a 7×9 tiled acquisition at 1386×1008 pixels per tile and 396 nm/pixel resolution and 13 z-planes per tile (axial resolution 1500 nm). Images were subjected to deconvolution to remove out-of-focus light. Acquired images were pre-processed (alignment and deconvolution with Microvolution software (http://www.microvolution.com/)) and segmented (including lateral bleed compensation) using publicly available CODEX image processing pipeline available at https://github.com/nolanlab/CODEX.

### Manual assessment of CELESTA performance on the HNSCC cohort

CELESTA performance on each sample of the HNSCC cohort was assessed manually by mapping CELESTA assigned cell types onto the CODEX images using the X and Y coordinates (**Figure S2**). For each cell type, CELESTA assigned cells were plotted as yellow crosses on the canonical marker staining images. Marker staining was shown as white signals on black background. Assessment for each cell were defined as positive canonical marker signals for that cell type. Manual assessment on CELESTA cell type assignments showed over 90% positive signals across the cell types on different samples.

### Manual gating of HNSCC cohort

The segmented CODEX HNSCC dataset was uploaded onto the Cytobank analysis platform and transformed with an inverse hyperbolic sine (asinh) transformation (cofactor of 5). The gating strategy used was as follows (**Figure S3**): Cells were defined by DRAQ5 nuclear expression and size, followed by endothelial cells (CD31+) and malignant cells (Cytokeratin+). CD4+ T cells (CD4+ CD8-CD3+ CD31-Cytokeratin-), CD8+ T cells (CD8+ CD4-CD3+ CD31-Cytokeratin-), *CD4/CD8 double positive cells (CD8+ CD4+ CD3+ CD31-Cytokeratin-), CD4/CD8 double negative cells (CD8- CD4- CD3+ CD31- Cytokeratin-),* and T regulatory cells (FOXP3+ CD25+ CD4+ CD8- CD3+ CD31- Cytokeratin-) were then defined. To adjust for the variability between sample image collection, each gate was tailored to each individual sample.

### Spatial analysis

We used co-location quotient (CLQ) ^23^ to identify cell spatial co-localization. If we define cell type *a* as target cells and cell type *b* as neighboring cells, the CLQ is the degree of cell type *b* co-locates with cell type *a* as ratio of the observed versus expected number of cell-type *b* among the set of nearest neighbors of cell-type *a* defined as:

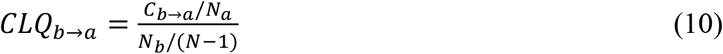

Where *C* is the number of cells of cell type *b* among the defined nearest neighbors of cell type *a. N* is the total number of cells and *N_a_* and *N_b_* are the numbers of cells for cell type *a* and cell type *b*. Cell types with fewer than twenty cells were excluded for each sample. We calculated a CLQ for every pair-wise cell types identified for each sample, and compared the CLQs on each pair of cell types between N+ and N0 samples to identify significantly different CLQs.

### Tumor tissue dissociation

Tumor tissue was thoroughly minced with a sterile scalpel and placed in a gentleMACS C-tube (Miltenyi Biotec, Sunnyvale, CA) containing 1.5mls of tissue digestion media. Tissue was mechanically digested on the GentleMACS dissociator five times under the human tumor tissue program h_tumor_01. Tissue was filtered with a 40μm nylon cell strainer (Falcon, Corning, NY) into a 14 ml tube filled up to 14 mls of tissue digestion media and spun at 4°C for 10 min at 514RCF. The mechanically digested cell pellet was re-suspended for 2 minutes on ice in 1-4 mls of ACK lysis buffer (Gibco, Grand Island, NY) depending on the pellet size and number of red blood cells present. Cells were filtered with a 40μm nylon cell strainer (Falcon, Corning, NY) into a 14 ml tube filled up to 14 mls of FACS buffer, Phosphate Buffered Saline without calcium or magnesium (Corning, Manassas, VA), 2%FBS heat inactivated, 10units/ml Penicillin-10ug/ml Streptomycin (Gibco, Grand Island, NY), 1mM Ultra Pure EDTA (Invitrogen, Carlsbad, CA) and spun at 4°C for 10 min at 514RCF. Cells were washed one more time with FACS buffer and re-suspended in 25μl of FACS buffer. Solid tissue in the strainer was collected and placed back in the C-tube with 2mls of tissue digestion media, 1ml of 3000U/ml collagenase/1000U/ml hyaluronidase (StemCell Technologies, Vancouver, BC) and 1 ml of 5U/ml dispase (StemCell Technologies, Vancouver, BC). The solid tissue in the C-tube was incubated at 37° Celsius on a rotator for 1 hour, then filtered with a 40μm nylon cell strainer (Falcon, Corning, NY) into a 14 ml tube filled up to 14 mls of tissue digestion media and spun at 4°C for 10 min at 514RCF. The enzymatically digested cell pellet was re-suspended in 1-4 mls of ACK lysis buffer (Gibco, Grand Island, NY) depending on the pellet size and number of red blood cells present for 2 minutes on ice. Cells were filtered with a 40μm nylon cell strainer (Falcon, Corning, NY) into a 14 ml tube filled up to 14 mls of FACS buffer, (Phosphate Buffered Saline without calcium or magnesium (Corning, Manassas, VA), 2%FBS heat inactivated, 10units/ml Penicillin-10ug/ml Streptomycin (Gibco, Grand Island, NY), 1mM Ultra pure EDTA (Invitrogen, Carlsbad, CA) and spun at 4°C for 10 min at 514RCF. Cells were re-suspended in FACS buffer, counted on a hemacytometer and washed one more time with FACS buffer. Cells were kept in FACS buffer on ice until flow cytometry staining. Sorting panel is shown in supplement **Table S5**.

### Single-cell RNA sequencing

RNA and library preparations were performed for 10x Genomics scRNA-sequencing samples according to 10x Genomics vs 2.0 handbook. Single cells were obtained according to the previous tumor tissue dissociation. Cells were stained with DAPI for live/dead detection and sorted for up to 500,000 live cells on a BD Aria II. Cells were counted after sort and right before 10x chip prep. 10x/Abseq by BD biosciences, followed the same protocol as the 10x Genomics samples except for the addition of FcBlock and Abseq antibody staining according to the manufacturer’s handbook.

### Singe-cell RNA-sequencing data processing and analysis

We aligned the reads using CellRanger. Preprocessing, data normalization and batch correction were done following Seurat SCTransform integration pipeline. UMAPs were obtained by using top 50 principal components. Cells were clustered by shared nearest neighbor modularity optimization. Cell types present were identified with canonical markers. Differentially expressed genes were identified in each cell type cluster between N0 and N+ patients using permutation test, and false discovery rate was used for multiple testing correction.

### Tissue microarray of HNSCC

Formalin-fixed paraffin-embedded tissue blocks of head and neck squamous cell carcinoma from 79 patients were pulled from the Stanford Health Care Department of Pathology archives. The area of malignancy was marked by a board-certified (C.S.K.) pathologist. Tissue microarrays were constructed from 0.6mm diameter cores punched from the tissue blocks. 4 um thick sections were stained with hematoxylin and eosin, FOXP3 (clone 236A/E7, 1:100 dilution; Leica BOND epitope retrieval solution 2) and cytokeratin mix (AE1/AE3, 1:75 dilution & CAM5.2, 1:25 dilution; Ventana Ultra; protease retrieval). The slides were digitized using Leica whole slide scanner with 40x magnification. Three samples with unknown lymph node status were excluded from analysis.

To assess co-localization of FOXP3 and cytokeratin immunohistochemistry staining, a preprocessing pipeline was built for the TMA data. The whole-slide images were dearrayed to obtain each core image. We next ran color deconvolution to quantify DAB staining using “scikit-image” package in Python. We thresholded the staining based on pixel intensity distributions of the DAB staining to quantify positive stained pixels in the images. We used a sliding window with 100 by 100 pixels to quantify the positive pixel densities for cytokeratin and FOXP3 within each window and move the sliding windows to cover the whole core area. We correlated the densities of cytokeratin staining and FOXP3 staining across sliding windows for each sample. We compared the density correlations between N0 and N+ samples.

### Statistical analysis and figure creation

Statistical analyses were performed and corresponding figures were generated in R or Python depending on the analysis packages. The student’s t-test, Wilcoxon rank-sum test and permutation test were utilized for comparisons between N0 and N+ samples. Specifically, for human patient data, student’s t-test was used for co-location quotient comparisons and density correlation comparisons between N0 and N+ sample. For *in vitro* and *in vivo* functional experiments, non-parametric Wilcoxon rank-sum test was used for comparisons. In addition, we created a paired-group setup in the transwell assay experiment and used paired test for statistical analysis. For multiple testing, permutation test was used and false discovery rate was used to adjust the p-values. Results were considered statistically significant when p < 0.05 or adjusted p<0.05 for multiple testing. Parts of Figure 5 and 6 were created using Biorender online tool (https://biorender.com). Multi-channel overlay images were created using ImageJ.

For the public colorectal cancer dataset, the “ground truth” is defined using the annotations in the publication ^6^. For the study cohort of HNSCC dataset, the “ground truth” standard is defined using manual gating based on gating strategies on cell types relevant to downstream spatial analysis. For each cell type, true positives (TP) is the number of cells assigned by both CELESTA and ground truth benchmark. False positives (FP) is the number of cells which were assigned by CELESTA but not ground truth benchmark. False negatives (FN) is the number of cells assigned in benchmark but not CELESTA. True negatives (TN) is the number of cells that are not assigned as the cell type by both CELESTA and benchmark. Precision is defined as TP/(TP+FP), and recall is defined as TP/(TP+FN). F1 score is defined as 2(precision*recall)/(precision+recall). Rand index to measure accuracy is defined as (TP + TN)/ (TP + TN + FP + FN). For the public colorectal cancer dataset, cell types with fewer than 5 cells in a sample region in the annotations were excluded. For HNSCC study cohort, adjusted rand index (ARI) was calculated for each sample using R package “mclust” adjustedRandIndex function.

### Cell lines and animals

B16-F0 parental and LN6-987AL murine melanoma cell lines have been described previously ^29^. Cells were cultured in Dulbecco’s Modified Eagle Medium (DMEM) supplemented with 4mM L-glutamine, 10% Fetal Bovine Serum (FBS), and 1% Penicillin Streptomycin. The tumor lines were routinely tested for mycoplasma by PCR, and all tests were negative. All animal studies were performed in accordance with the Stanford University Institutional Animal Care and Use Committee under protocol APLAC-17466. All mice were housed in an American Association for the Accreditation of Laboratory Animal Care-accredited animal facility and maintained in specific pathogen-free conditions.

### Transwell migration assays

FoxP3^EGFP^ mice ^42^ were acquired from Jackson (006772) and bred in our facility at Stanford University. Splenocytes were harvested from tumor-naive female FoxP3^EGFP^ mice. Spleens were subjected to mechanical dissociation on 70μm cell strainers and washed with HBSS supplemented with 2% FBS and 2mM Ethylenediaminetetraacetic acid (EDTA) (HBSSFE). Erythrocytes were lysed with Ammonium-Chloride-Potassium (ACK). Magnetic isolation of Tregs was performed using the EasySep Mouse CD25 Regulatory T cell Positive Selection Kit (StemCell, 18782) according to the manufacturer’s instructions. Tregs were cultured in RPMI-1640 supplemented with 10% FBS, 2mM L-glutamine, 15mM HEPES, 14.3 mM 2-mercaptoethanol, 1mM Sodium Pyruvate, 1 × MEM Non-Essential Amino Acids Solution, and 300IU hIL-2 (Peprotech) for 72 hours.

Tumor cell line suspensions were prepared by washing with phosphate buffered saline (PBS) followed by treatment with StemPro Accutase (Thermo, A1110501). 10^5^ tumor cells were plated in the bottom chamber of the 24-well transwell plates 24 hours prior to the assay. 5μm transwell membranes (Costar, 3421) were incubated in complete RPMI for 24 hours prior to the assay. Membranes were transferred to the tumor-containing wells and suspensions of 5×10^4^ Tregs were added to the top chambers of the transwells. Cells were cultured for 2 hours at 37°C in 5% CO_2_, after which the membranes were removed, and cells from the bottom chamber were processed for analysis by flow cytometry.

Cell suspensions were washed in HBSSFE and stained with the following antibodies: Mouse Fc Block (BD, 2.4G2), CD4 (BioLegend, RM4-5), CD25 (BioLegend, PC61), and CXCR3 (BioLegend, CXCR3-173). 4’,6-Diamidino-2-Phenylindole Dihydrochloride (DAPI) was used to stain for viability. Samples were run on an LSRFortessa cytometer (Becton Dickinson) and analyzed using FlowJo V10 software (TreeStar).

### In vivo Treg tumor infiltration

Experiments were performed using either C57NL/6J (Jackson, 000664) or FoxP3^EGFP^ (Jackson, 006772) female mice housed in our facility at Stanford. B16-F0 or LN6-987AL tumor cells were washed with PBS and dissociated from tissue culture plastic with StemPro Accutase (Thermo, A1110501). Cell suspensions of 2×10^5^ cells in phenol-red free DMEM were injected into the subcutaneous region of the left flank of nine-week-old female mice (Jackson, 000664) following removal of fur with surgical clippers. After 15 days of tumor growth, mice were euthanized and their tumors were processed for analysis by flow cytometry.

Tumors were weighed followed by digestion in RPMI-1640 supplemented with 4mg/mL Collagenase Type 4 (Worthington, LS004188) and 0.1mg/mL Deoxyribonuclease I (DNAse I, Sigma, DN25) at 37°C for 20 minutes with agitation. Tumors were then dissociated on 70μm strainers, washed with HBSSFE, and stained for viability using LIVE/DEAD Fixable Blue Dead Cell Stain (Thermo, L34962). Surface proteins were stained, samples were fixed and permeabilized using the eBioscience FoxP3 Fixation/Permeabilization kit (Thermo, 00-5521-00), and intracellular FoxP3 was stained. The following antibodies were used: Mouse Fc Block (BD, 2.4G2), CD4 (BioLegend, RM4-5), CD8α (BioLegend, 53-6.7), CD3 (BioLegend, 17A2), CD25 (BioLegend, PC61), CXCR3 (BioLegend, CXCR3-173), B220 (BioLegend, RA3-6B2), CD45.2 (BioLegend, 104), and FoxP3 (Thermo/eBiosciences, NRRF-30). AccuCount Fluorescent particles (Spherotec, ACFP-50-5) were added to each samples for the purposes of determining absolute cell counts. Samples were run on an LSRFortessa cytometer (Becton Dickinson) and analyzed using FlowJo V10 software (TreeStar).

For treatment CXCR3-blockade studies, LN6-987AL cells were prepared as above and injected into seven-week-old FoxP3^EGFP^ mice. Mice were treated with AMG487 (R&D Systems, 4487) at 5mg/kg every 48hrs starting on day one following tumor implantation. After 9 days of tumor growth, mice were euthanized and their tumors were processed for analysis by flow cytometry as described above.

## ACKNOWLEDGEMENT

This work was supported by the National Institute of Health, National Cancer Institute U54 CA209971.

## DATA AND CODE AVAILABILITY

The scRNA-seq data are deposited at GEO: GSE140042. HNSCC imaging data are hosted at Synapse.org SageBionetworks at https://doi.org/10.7303/syn26242593. The benchmark public imaging data can be found at https://doi.org/10.7937/tcia.2020.fqn0-0326. All codes related to CELESTA can be found at https://github.com/plevritis/CELESTA.

