## Supplemental Figures 1-6 and supplemental Tables 1-5 for "Identification of cell types in multiplexed *in situ* images by combining protein expression and spatial information using CELESTA reveals novel spatial biology"

### SUPPLEMENT

#### *Supplement figures*

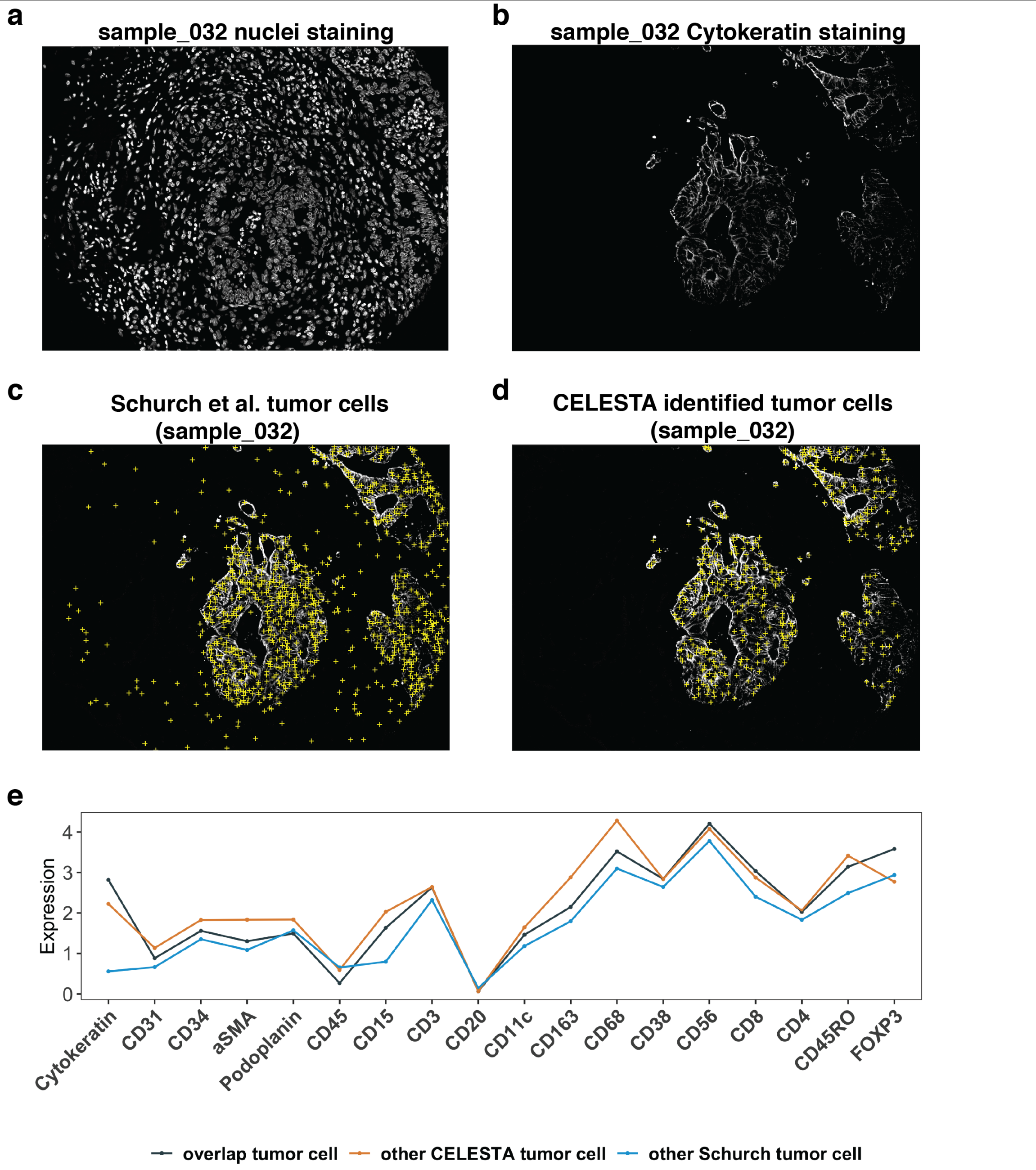


**Figure S1. Example illustrating differences on tumor cell identification between CELESTA and Schurch et al. annotations on the colorectal cancer dataset. (a)** Nuclei staining of one example core region (032). **(b)** Cytokeratin staining of region 032. **(c)** Tumor cells identified in Schurch et al. annotations of region 032 (yellow crosses) overlaid on cytokeratin staining. **(d)** Tumor cells identified by CELESTA of region 032 (yellow crosses) overlaid on cytokeratin staining. **(e)** Average canonical cell type marker expressions across all the 70 samples on cells both CELESTA and Schurch et al. identified to be tumor cells (overlap tumor cells, black), other cells only CELESTA identified to be tumors (orange) and other tumor cells in the Schurch et al. tumor cluster (blue). Tumor cells identified by CELESTA have to express cytokeratin.

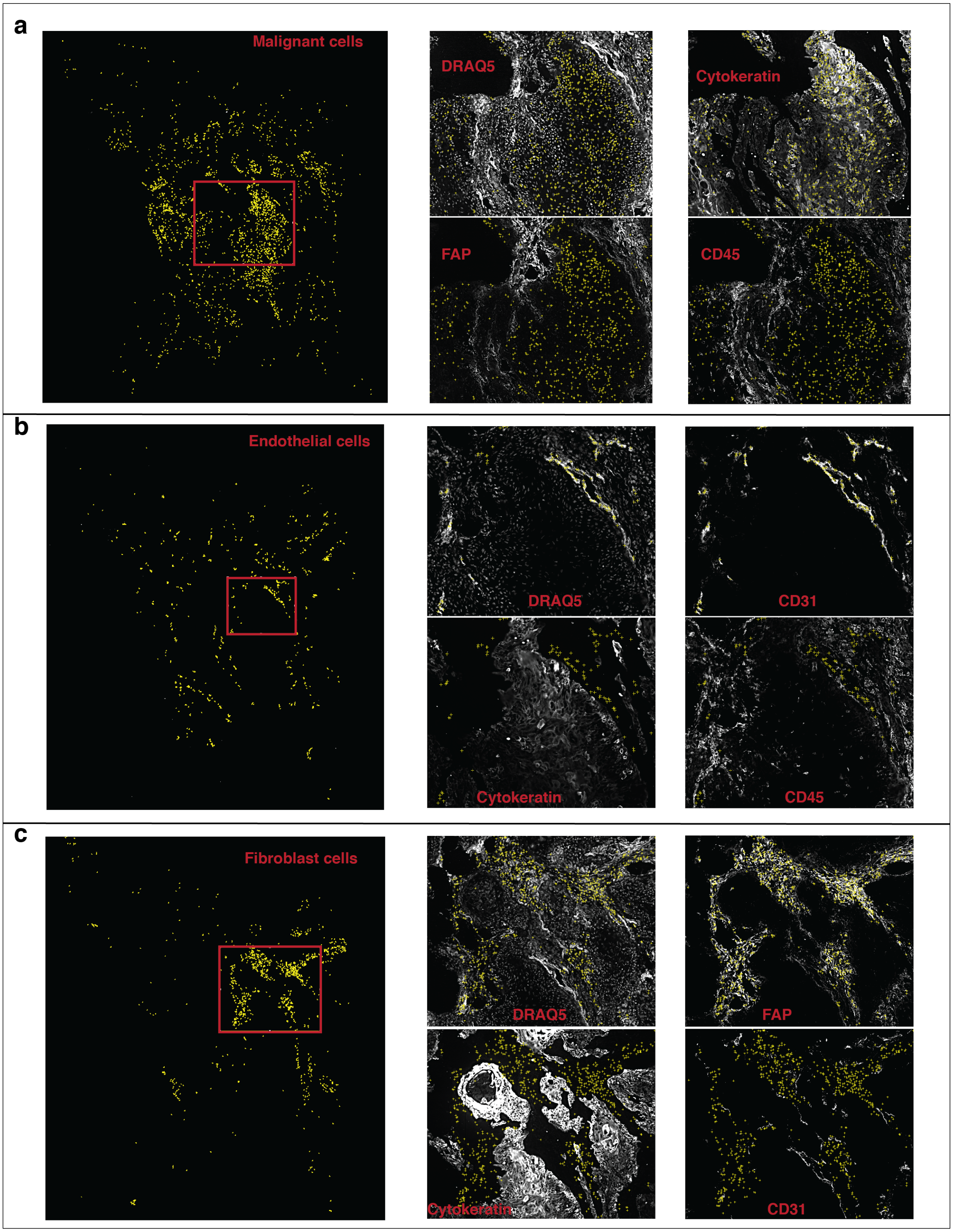

a
Malignant cells
DRAQ5
Cytokeratin
FAP
CD45
b
Endothelial cells
DRAQ5
CD31
Cytokeratin
CD45
c
Fibroblast cells
DRAQ5
FAP
Cytokeratin
CD31

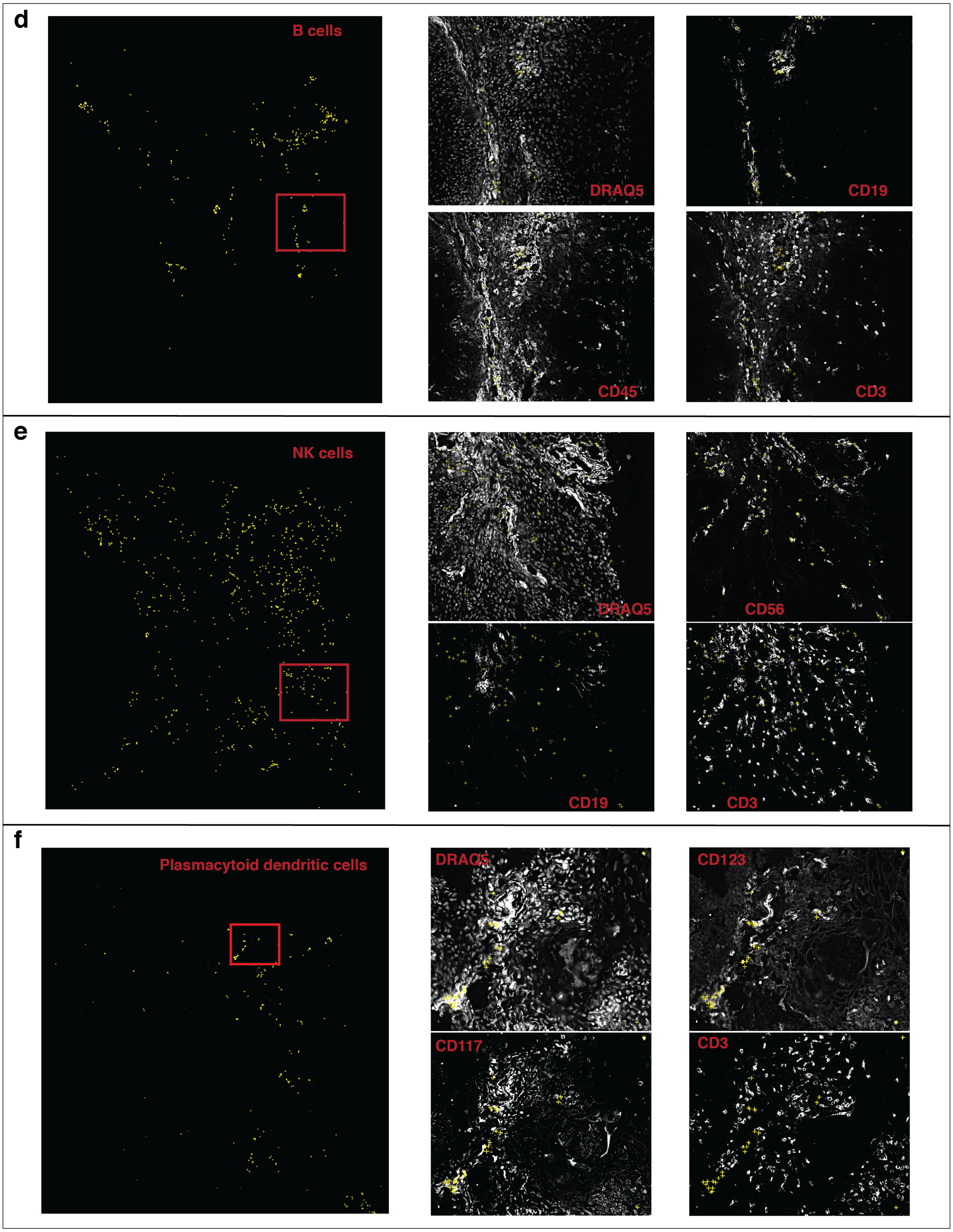
d
B cells
DRAQ5
CD19
CD45
CD3
e
NK cells
DRAQ5
CD56
CD19
CD3
f
Plasmacytoid dendritic cells
DRAQ5
CD123
CD117
CD3

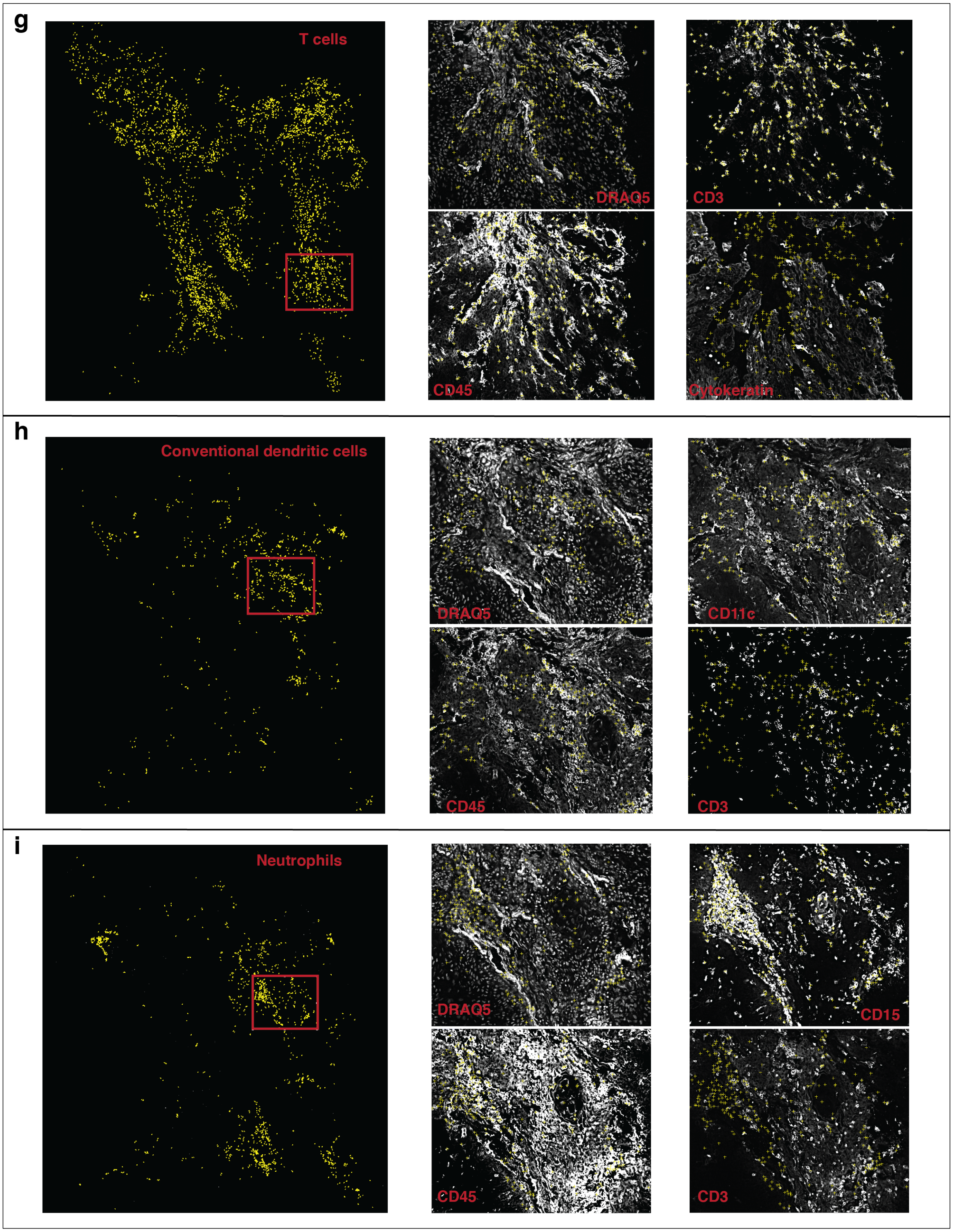
g
T cells
DRAQ5
CD3
CD45
Cytokeratin
h
Conventional dendritic cells
DRAQ5
CD11c
CD45
CD3
i
Neutrophils
DRAQ5
CD15
CD45
CD3

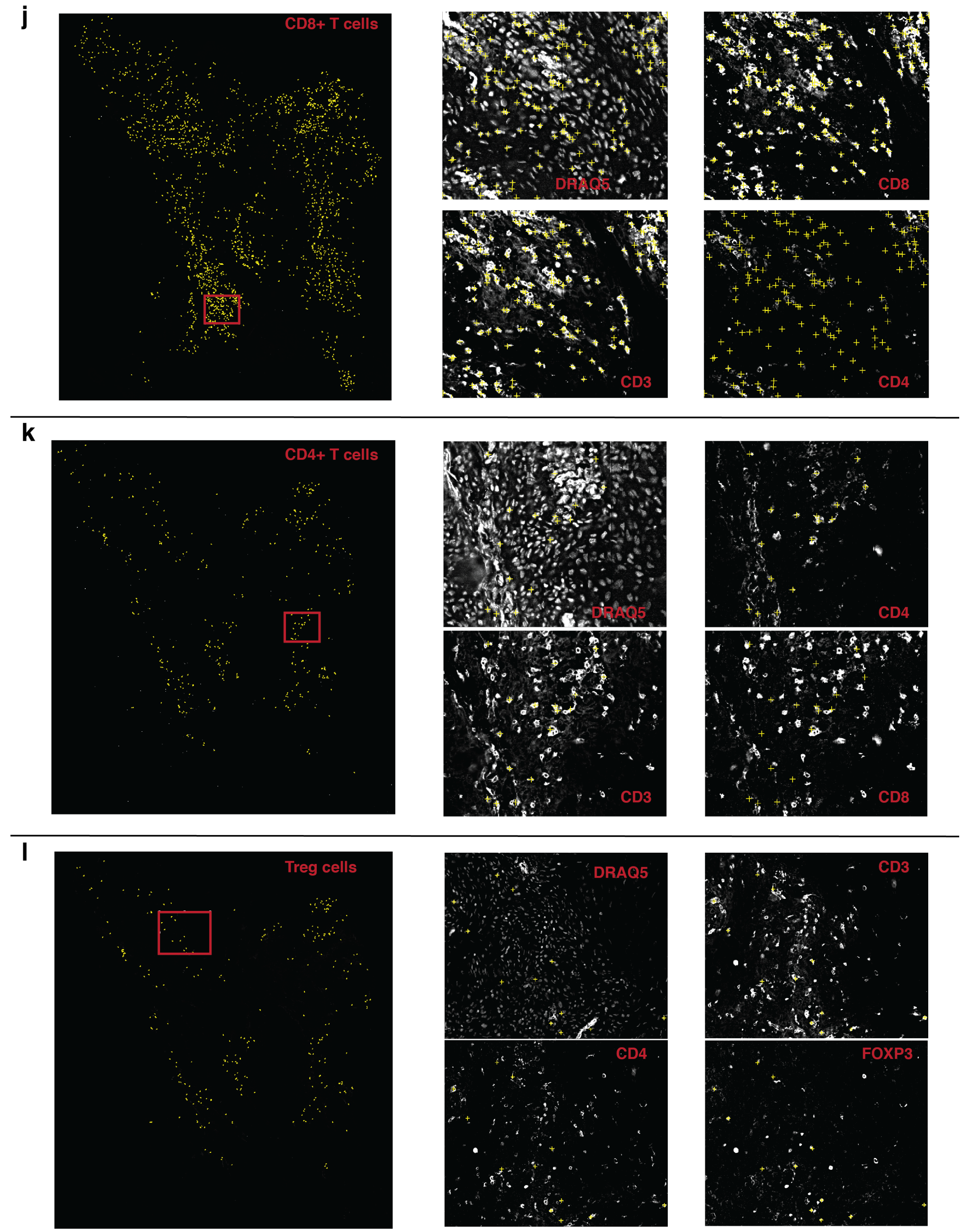


**Figure S2. Manual assessment of CELESTA identified cell types on an example HNSCC sample. (a)-(l)** Identified cells are shown as yellow crosses using the X and Y coordinates overlaid on canonical marker staining (white) CODEX images. For each cell type, nuclei staining and three example markers (positive and negative) important for the cell type are shown.

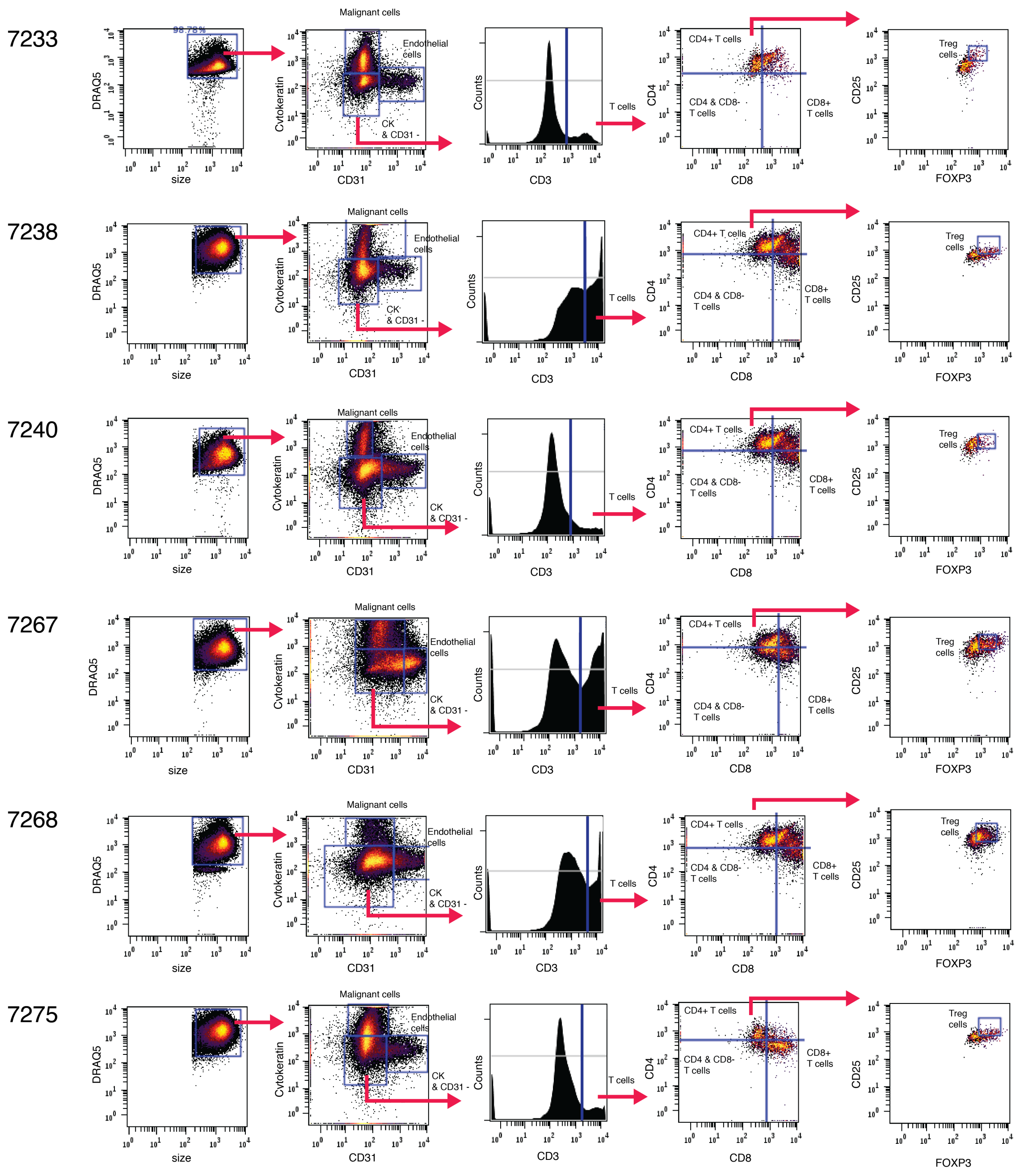


**Figure S3. Gating strategies on the head and neck squamous cell carcinoma (HNSCC) samples.** Gating strategies used to identify key cell types relevant to the HNSCC study including malignant cells, endothelial cells and subtypes of T cells.

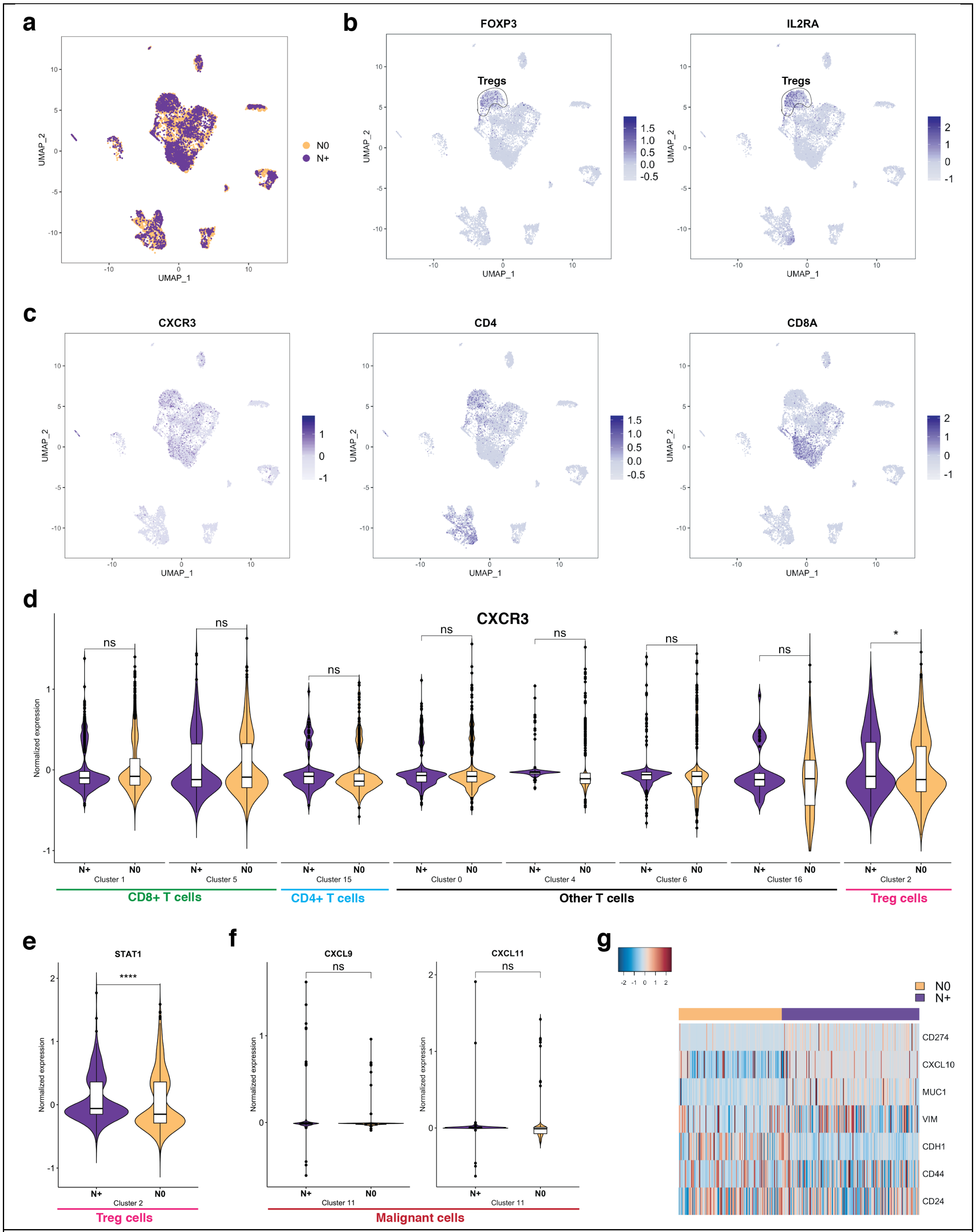


**Figure S4. Additional scRNA-seq analysis. (a)** UMAP plot of identified cell type clusters with node status. **(b)-(c)** UMAP plots of FOXP3, IL2RA, CXCR3, CD4 and CD8A. **(d)** CXCR3 expression in different T cell clusters showed that CXCR3 only differentially expressed in Treg cells. **(e)** Violin plot of STAT1 expression in the Treg cell cluster between N+ and N0 samples. STAT1 is a CXCR3 inducer. **(f)** Violin plot of CXCL9 and CXCL11 in the malignant cell cluster between N+ and N0 samples. CXCL9 and CXCL11 are both ligands of CXCR3, but they are not differentially expressed in our data. **(g)** Heatmap shows expressions of CD274 (PD-L1), MUC1, EMT markers (CDH1 and VIM) and stemness markers (CD44 and CD24). This observation suggests that malignant cells in the N0 patients are more in the epithelial state while malignant cells in the N+ patients are more mesenchymal, consistent with the fact that mesenchymal state cells are easier to migrate and related to poor prognosis. *: adjusted p-value < 0.05, **: adjusted p-value < 0.01, ***: adjusted p-value < 0.005, ****: adjusted p-value < 0.001.

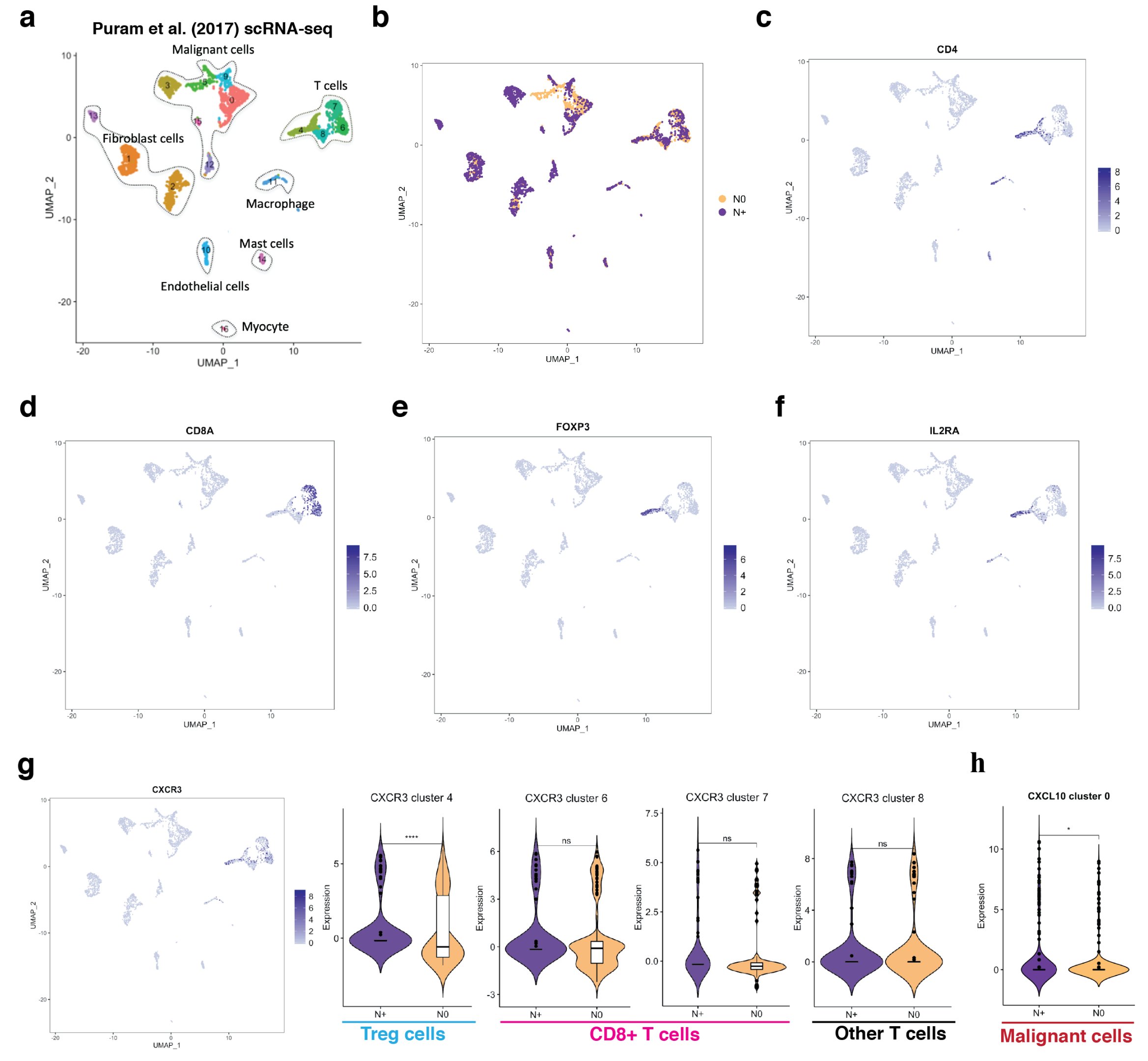


**Figure S5. Additional scRNA-seq analysis using public domain data from Puram et al. (2017). (a)** UMAP plot of identified cell type clusters. **(b)** UMAP plot of identified cell type clusters with node status. **(c)-(f)** UMAP plots of CD4, CD8A, FOXP3, and IL2RA. **(g)** UMAP plot of CXCR3 and violin plots of CXCR3 in the T cell clusters. **(h)** Violin plot of CXCL10 in malignant cell cluster 0. *: adjusted p-value < 0.05, **: adjusted p-value < 0.01, ***: adjusted p-value < 0.005, ****: adjusted p-value < 0.001.

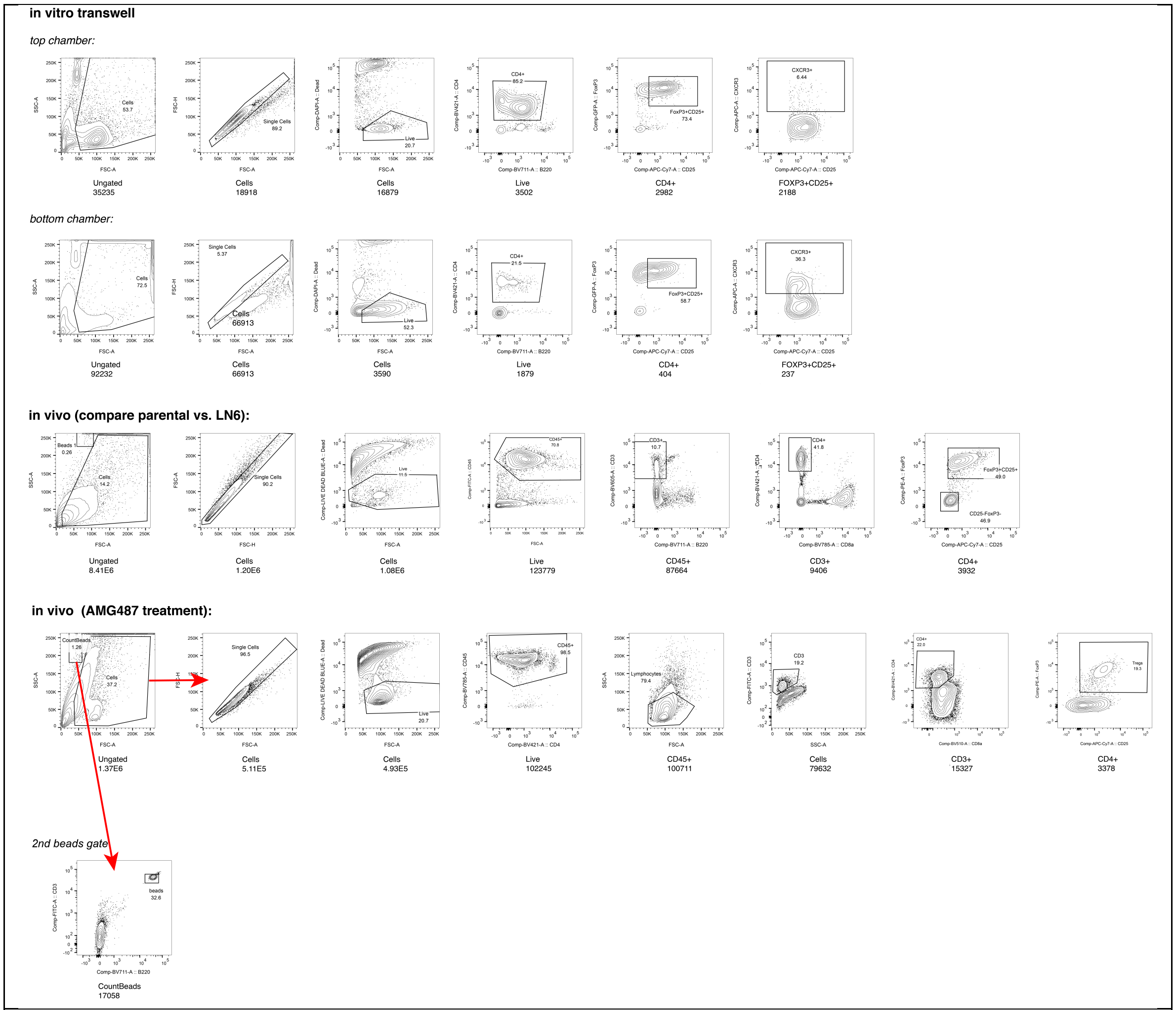


**Figure S6. Example gating strategies used for mouse model studies.**

#### *Supplement tables*

| Cell type | CD31 | CD34 | CK | aSMA | PDPN | CD45 | CD15 | CD3 | CD20 | CD11c | CD163 | CD68 | CD38 | CD56 | CD8 | CD4 | CD45RO | FOXP3 | Round |
|---|---|---|---|---|---|---|---|---|---|---|---|---|---|---|---|---|---|---|---|
| **Endothelial** | 1 | 1 | 0 | 0 | 0 | 0 | 0 | 0 | 0 | 0 | 0 | 0 | 0 | 0 | 0 | 0 | 0 | 0 | 1 |
| **Malignant** | 0 | 0 | 1 | 0 | 0 | 0 | 0 | 0 | 0 | 0 | 0 | 0 | 0 | 0 | 0 | 0 | 0 | 0 | 1 |
| **aSMA+ stroma** | 0 | 0 | 0 | 1 | 0 | 0 | 0 | 0 | 0 | 0 | 0 | 0 | 0 | 0 | 0 | 0 | 0 | 0 | 1 |
| **Lymphatics** | 0 | 0 | 0 | 0 | 1 | 0 | 0 | 0 | 0 | 0 | 0 | 0 | 0 | 0 | 0 | 0 | 0 | 0 | 1 |
| **Immune (general)** | 0 | 0 | 0 | 0 | 0 | 1 | NA | NA | NA | NA | NA | NA | NA | NA | NA | NA | NA | NA | 1 |
| **T cell** | 0 | 0 | 0 | 0 | 0 | NA | 0 | 1 | 0 | 0 | 0 | 0 | 0 | 0 | NA | NA | NA | NA | 2 |
| **Granulocyte** | 0 | 0 | 0 | 0 | 0 | NA | 1 | 0 | 0 | 0 | 0 | NA | 0 | 0 | 0 | 0 | NA | 0 | 2 |
| **B cell** | 0 | 0 | 0 | 0 | 0 | NA | 0 | 0 | 1 | 0 | 0 | 0 | 0 | 0 | 0 | 0 | 0 | 0 | 2 |
| **CD11c+ DC** | 0 | 0 | 0 | 0 | 0 | NA | 0 | 0 | 0 | 1 | 0 | 0 | 0 | 0 | 0 | 0 | 0 | 0 | 2 |
| **CD68+CD163+ Macrophage** | 0 | 0 | 0 | 0 | 0 | NA | 0 | 0 | 0 | 0 | 1 | 1 | 0 | 0 | 0 | 0 | NA | 0 | 2 |
| **Plasma cell** | 0 | 0 | 0 | 0 | 0 | NA | 0 | 0 | 0 | 0 | 0 | 0 | 1 | 0 | 0 | 0 | 0 | 0 | 2 |
| **NK cell** | 0 | 0 | 0 | 0 | 0 | NA | 0 | 0 | 0 | 0 | 0 | 0 | 0 | 1 | 0 | 0 | 0 | 0 | 2 |
| **CD8+ T cell** | 0 | 0 | 0 | 0 | 0 | NA | 0 | NA | 0 | 0 | 0 | 0 | 0 | 0 | 1 | 0 | NA | 0 | 3 |
| **CD4+ T cell** | 0 | 0 | 0 | 0 | 0 | NA | 0 | NA | 0 | 0 | 0 | 0 | 0 | 0 | 0 | 1 | NA | NA | 3 |
| **CD4+CD45RO+ T cell** | 0 | 0 | 0 | 0 | 0 | NA | 0 | NA | 0 | 0 | 0 | 0 | 0 | 0 | 0 | 1 | 1 | 0 | 4 |
| **Treg cell** | 0 | 0 | 0 | 0 | 0 | NA | 0 | NA | 0 | 0 | 0 | 0 | 0 | 0 | 0 | 1 | 0 | 1 | 4 |

**Table S1:** An example of the initial cell-type signature matrix used in CELESTA based on the CODEX panel used for the colorectal cancer FFPE samples (Schurch et al. 2020). DC: dendritic cell. CK: cytokeratin. PDPN: podoplanin.

| Patient ID | Stage | Dataset |
|---|---|---|
| **SCC7153** | T3/N0/Mx | CODEX |
| **SCC7155** | T4a/N0/Mx | CODEX |
| **SCC7233** | T3/N3b/Mx | CODEX |
| **SCC7238** | T4a/N2b/Mx | CODEX & scRNA-seq |
| **SCC7240** | T3/N2b/Mx | CODEX & scRNA-seq |
| **SCC7267** | T4a/N3/Mx | CODEX |
| **SCC7268** | T3/N0/Mx | CODEX & scRNA-seq |
| **SCC7275** | T4a/ N0/Mx | CODEX & scRNA-seq |

**Table S2:** Staging information of head and neck squamous cell carcinoma (HNSCC) samples included in the study.

**Immune markers**

CD19 CD31 CD7 CD15 CD21 CD3
CD21 CD4 CD69 CD2 CD25 CD66 CD45
CD8 CD11c CD56 CD38 CD36 CD9
CD1c CD123 Foxp3 CD40 CD117 CD5

**Functional markers**

LILRB1 HLA-DR Ki67 IgM PD1
PD-L1 CD152 CD57 CD16 ICOS
HIF-1a CD154 CD54 CD127
CD49f MMP12 TCRyd CD185

**Auxiliary markers**

Vimentin FAP CD90 CD45RA
Podoplanin Collagen IV CD273
Cytokeratin HLA-ABC SLC2A1

**Table S3:** OCT CODEX panel of head and neck squamous cell carcinoma study.

| Cell type | CD45 | CD31 | CK | FAP | CD3 | CD4 | CD19 | CD56 | CD8 | Foxp3 | CD25 | CD11c | CD117 | CD15 | CD123 | CD66 | Round |
|---|---|---|---|---|---|---|---|---|---|---|---|---|---|---|---|---|---|
| **Endothelial** | 0 | 1 | 0 | 0 | 0 | 0 | 0 | 0 | 0 | 0 | 0 | 0 | 0 | 0 | 0 | 0 | 1 |
| **Malignant** | 0 | 0 | 1 | 0 | 0 | 0 | 0 | 0 | 0 | 0 | 0 | 0 | 0 | 0 | 0 | 0 | 1 |
| **Fibroblast** | 0 | 0 | 0 | 1 | 0 | 0 | 0 | 0 | 0 | 0 | 0 | 0 | 0 | 0 | 0 | 0 | 1 |
| **Immune(general)** | 1 | 0 | 0 | 0 | NA | NA | NA | NA | NA | NA | NA | NA | NA | NA | NA | NA | 1 |
| **Myeloid** | 1 | 0 | 0 | 0 | 0 | 0 | 0 | 0 | 0 | 0 | 0 | NA | 1 | NA | NA | 1 | 2 |
| **T cell** | 1 | 0 | 0 | 0 | 1 | NA | 0 | 0 | NA | NA | NA | 0 | 0 | 0 | 0 | 0 | 2 |
| **B cell** | 1 | 0 | 0 | 0 | 0 | 0 | 1 | 0 | 0 | 0 | 0 | 0 | 0 | 0 | 0 | 0 | 2 |
| **NK cell** | 1 | 0 | 0 | 0 | 0 | 0 | 0 | 1 | 0 | 0 | 0 | 0 | 0 | 0 | 0 | 0 | 2 |
| **Neutrophil** | 1 | 0 | 0 | 0 | 0 | 0 | 0 | 0 | 0 | 0 | 0 | 0 | 0 | 1 | 0 | NA | 3 |
| **pDC** | 1 | 0 | 0 | 0 | 0 | 0 | 0 | 0 | 0 | 0 | 0 | 0 | 1 | 0 | 1 | NA | 3 |
| **cDC** | 1 | 0 | 0 | 0 | 0 | 0 | 0 | 0 | 0 | 0 | 0 | 1 | 1 | 0 | 0 | NA | 3 |
| **CD4+ T cell** | 1 | 0 | 0 | 0 | 1 | 1 | 0 | 0 | 0 | NA | NA | 0 | 0 | 0 | 0 | 0 | 4 |
| **CD8+ T cell** | 1 | 0 | 0 | 0 | 1 | 0 | 0 | 0 | 1 | 0 | 0 | 0 | 0 | 0 | 0 | 0 | 4 |
| **Treg cell** | 1 | 0 | 0 | 0 | 1 | 1 | 0 | 0 | 0 | 1 | 1 | 0 | 0 | 0 | 0 | 0 | 5 |

**Table S4:** An example of initial cell-type signature matrix used in CELESTA based on the OCT CODEX panel in the head and neck squamous cell carcinoma study. pDC: plasmacytoid dendritic cell. cDC: conventional dendritic cell. CK: cytokeratin.

| **Category** | **Fluorescence** | **Vendor** | **Marker** | **Clone** | **Catalog #** | **Lot #** |
|---|---|---|---|---|---|---|
| Immune | FITC | Biolegend | CD45 | 2D1 | 368508 | B230036 |
| Immune | FITC | Biolegend | CD19 | HIB19 | 302206 | B248329 |
| Immune | FITC | Biolegend | CD68 | Y1/82A | 333806 | B243370 |
| Immune | FITC | Biolegend | CD3 | HIT3a | 300306 | B218086 |
| Endothelial | PE | BD | CD31 | WM59 | 555446 | 7335512 |
| Endothelial | PE | Biolegend | CD140a(PDGFRa) | 16A1 | 323506 | B233466 |
| Fibroblasts | APC | Biolegend | FAP | 427819 | FAB3715A | AEH10116051 |
| Live | DAPI | Biolegend | LIVE/DEAD | | 422801 | |

**Table S5:** Sorting panel.
